# A role for the Smc3 hinge domain in the maintenance of sister chromatid cohesion

**DOI:** 10.1101/179788

**Authors:** Brett Robison, Vincent Guacci, Douglas Koshland

**Affiliations:** Department of Molecular and Cell Biology, University of California, Berkeley Berkeley, CA 94720

**Keywords:** sister chromatid cohesion, condensation, cohesin, Smc3, hinge, Pds5, Eco1

## Abstract

Cohesin is a conserved protein complex required for sister chromatid cohesion, chromosome condensation, DNA damage repair, and regulation of transcription. Although cohesin functions to tether DNA duplexes, the contribution of its individual domains to this activity remains poorly understood. We interrogated the Smc3p subunit of cohesin by random insertion mutagenesis. Analysis of a mutant in the Smc3p hinge revealed an unexpected role for this domain in cohesion maintenance and condensation. Further investigation revealed that the Smc3p hinge functions at a step following cohesin’s stable binding to chromosomes and independently of Smc3p’s regulation by the Eco1p acetyltransferase. Hinge mutant phenotypes resemble loss of Pds5p, which binds opposite the hinge near Smc3p’s head domain. We propose that a specific conformation of the Smc3p hinge and Pds5p cooperate to promote cohesion maintenance and condensation.

## Introduction

Cohesin is a conserved protein complex required for sister chromatid cohesion, chromosome condensation, DNA damage repair, and regulation of transcription (Onn et al. 2008). To accomplish these functions, chromosome-bound cohesin tethers two distinct DNA duplexes or two sites on a single DNA duplex. A remarkable feature of cohesin-mediated tethers is that they must persist for long periods. For example, once generated, cohesion between sister chromatids must be maintained for up to several hours until cells progress through mitosis. Cohesion maintenance is essential for a successful mitosis since it ensures bipolar attachment and proper segregation of chromosomes. This process is crucial in mammalian oocytes since cohesion must be maintained from its establishment during meiotic prophase I, which occurs during fetal development, until the egg is fertilized in adulthood. Failure to maintain this cohesion can lead to aneuploidy and may cause infertility and birth defects in humans (Duncan et al. 2012). However, despite its critical function, the mechanism and regulation of cohesion maintenance remains poorly understood.

Cohesin is a large multi-subunit complex with an elaborate molecular architecture. In the budding yeast *Saccharomyces cerevisiae*, core cohesin subunits are Smc1p, Smc3p, Mcd1p (also called Scc1p), and Scc3p (Onn et al. 2008). The Structural Maintenance of Chromosome (Smc) proteins fold back on themselves to form large dumbbell-shaped structures with two globular domains, referred to as the head and hinge, separated by an ∼45 nm long coiled coil (Onn et al. 2008). Cohesin or purified Smc1p-Smc3p heterodimers have been visualized by electron microscopy, atomic-force microscopy, and scanning-force microscopy (Haering et al. 2002; Sakai et al. 2003; Kulemzina et al. 2016). These studies revealed that Smc1p and Smc3p dimerize by an interaction between their heads and a separate interaction between their hinges. Dimerization of the heads is further stabilized by the kleisin subunit Mcd1p which binds through its N-terminus to Smc3p and its C-terminus to Smc1p (Haering et al. 2002). The existence of two dimerization interfaces allows cohesin to form large rings. This ring structure likely explains cohesin’s ability to bind DNA by topological entrapment. In addition to these ring structures, more complex conformations have also been observed (Sakai et al. 2003). Evidence supporting the biological significance of these other conformations has been lacking.

Sister chromatid cohesion is established in S phase then maintained until anaphase onset. Cohesion establishment is a multi-step process. In budding yeast, the Scc2p/Scc4p complex (Ciosk et al. 2000) loads cohesin onto DNA at centromeres and along chromosome arms at <u>c</u>ohesin-<u>a</u>ssociated regions or CARs in early S phase (Megee et al. 1999; Laloraya et al. 2000; Glynn et al. 2004). During S phase, DNA-bound cohesin is converted into a form that tethers sister chromatids by the Eco1p acetyltransferase, which acetylates Smc3p at lysines 112 and 113 (Toth et al. 1999; Skibbens et al. 1999; Ünal et al. 2008; Ben-Shahar et al. 2008; Zhang et al. 2008). Once cohesion is established in S phase, the cohesion-associated regulator Pds5p is required to maintain cohesion until anaphase onset (Hartman et al. 2000; Panizza et al. 2000; Stead et al. 2003).

The mechanism of cohesion maintenance is only partially understood. Pds5p co-localizes with cohesin on chromosomes and when mutated, causes a decrease in cohesin binding to chromosomes, a reduction in cellular Mcd1p levels, and a cohesion maintenance defect (Hartman et al. 2000; Panizza et al. 2000). This maintenance defect can be suppressed by preventing premature Mcd1p degradation via a polySUMO-dependent pathway, or preserving Smc3p acetylation by deleting the *HOS1* deacetylase (Stead et al. 2003; D’Ambrosio et al. 2014; Chan et al. 2013). Thus, Pds5p may function to protect cohesin complex from factors that could dissolve cohesion. However, cohesion maintenance is a more complex process. The cohesin mutant Mcd1-ROCC is defective for cohesion maintenance yet Mcd1p levels are not reduced and Pds5p recruitment to cohesin and chromosomes is unaffected (Eng et al. 2014). These observations suggest that an additional step beyond Mcd1p stabilization or Pds5p recruitment is required for cohesion maintenance.

A clue for this additional step comes from imaging and biochemical studies of cohesin and Pds5p. Biochemical studies indicate Pds5p binds to Mcd1p, placing Pds5p adjacent to the Smc head domains (Chan et al. 2013; Lee et al. 2016; Muir et al. 2016; Ouyang et al. 2016). The functional significance of this interaction is supported by mutations in budding yeast Mcd1p that mimic the cohesion maintenance defects upon Pds5p depletion (Eng et al. 2014). However, crosslinking has shown human Pds5Bp interacts with all cohesin subunits, implying that its association with cohesin is more extensive and/or dynamic (Huis in t Veld et al. 2014; Hons et al. 2016). Furthermore, *in vivo* FRET suggested that Pds5p localizes near the hinge (Mc Intyre et al. 2007) and atomic force microscopy shows Smc1p/Smc3p dimer conformations in which the hinge and head regions are adjacent (Sakai et al. 2003). This proximity was supported by the observation that purified hinge domains are capable of binding to the head-associated Scc3p subunit of cohesin (Murayama and Uhlmann 2015). Scc3p binds to the head and also binds Pds5p. Taken together these biochemical results suggest that cohesion might be maintained by an unanticipated conformation of cohesin involving binding of the hinge to the head.

Given the evidence that Pds5p has interactions with both the head and hinge regions, it is unclear how Pds5p mediates cohesion maintenance and which Smc domains are involved. To begin to address these issues, we conducted a comprehensive RID screen of Smc3p, a transposon-based mutagenesis approach that generates random 5 amino acid insertions. Here we characterize an insertion mutant located in the Smc3p hinge region. This mutant establishes cohesion but fails to maintain it, yet Pds5p remains bound to cohesin and to chromosomes. Previous work suggested that the Smc hinge region functions only in cohesion establishment (Gruber et al. 2006; Kurze et al. 2011). Our analysis reveals that the Smc3p hinge is important for cohesion maintenance.

## Results

### The D667 region of the Smc3p hinge enhances but is not essential for cohesin binding at centromeres and cohesin-associated regions

We used a random insertion dominant (RID) screen to identify partial loss of function alleles of *SMC3* (Milutinovich et al. 2007; Eng et al. 2014). We expected to obtain RID screen mutations at the interfaces between Smc3p and Smc1p or Mcd1p. These mutations would be expected to prevent assembly and subsequent loading of cohesin onto chromosomes. In addition to assembly mutants, we predicted that mutations that preserved cohesin assembly would be found. We reasoned that if Smc3p function is modulated after cohesin assembles and binds chromosomes to maintain cohesion, mutants of Smc3p could be found that impair this step.

Mutant *SMC3* alleles were generated by *in vitro* transposon-mediated mutagenesis, which produced a library encoding random five-amino acid insertions (Supplemental Figure 1, Materials and Methods). In this library, *SMC3* was placed under control of the conditional *pGAL1* promoter. The library was transformed into both wild-type haploid yeast and the temperature-sensitive *smc3-42* strain. Transformants were obtained on dextrose-containing media to repress RID library *pGAL1-SMC3* expression. Colonies were then screened for impaired growth on plates containing galactose as the carbon source to drive *pGAL1*-mediated overexpression of mutant *SMC3* alleles. The location of insertions within *SMC3* that impaired growth of wild-type (Supplemental Table 1) or *smc3-42* cells (Supplemental Table 2) when overexpressed were then determined by sequencing.

**Figure 1:**
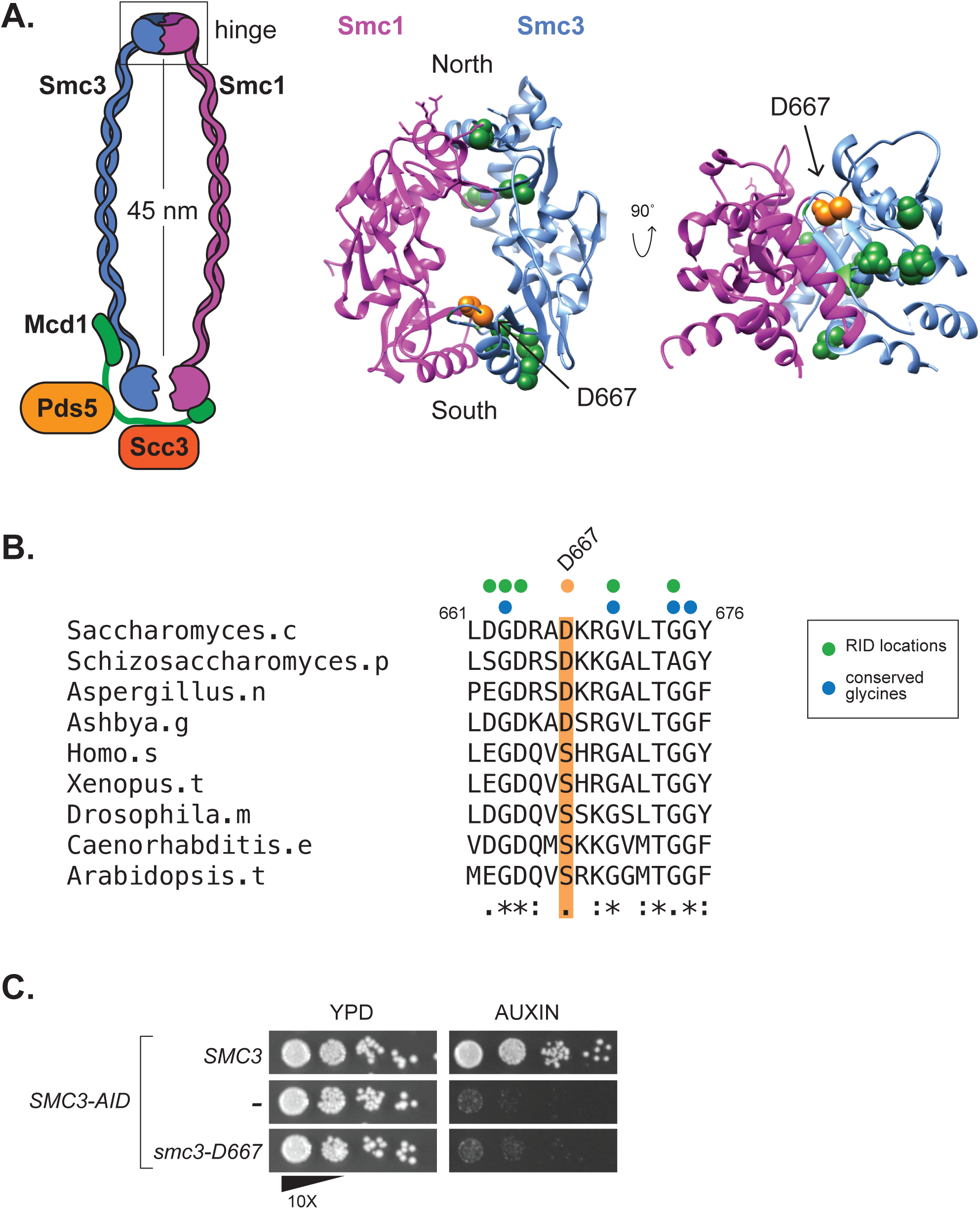
The *smc3-D667* RID mutation maps to a loop near the South interface of the Smc3p hinge. A. Diagram of cohesin highlighting location of the *smc3-D667* RID insertion. The homologous residue of *smc3-*D667, highlighted in orange, was determined by sequence alignment using ClustalW and mapped onto the mouse Smc1p/Smc3p hinge crystal structure (PDB: 2WD5, Kurze et al. 2011). Other RIDs isolated in this screen and located in the hinge domain are represented as green spheres and their positions were also approximated by sequence alignment. B. Sequence alignment of Smc3p homologues showing the conserved region around D667. The position of Asp667 is highlighted in orange and the sequence of the five-amino acid insertion, AAAAD, that follows Asp667 in the *smc3-D667* RID is depicted above as an orange dot. The position of other RIDs in this region are shown with green dots, and conserved glycine residues shown with blue dots. C. The *smc3-D667* allele under the native *SMC3* promoter is unable to support viability. Cultures of haploid strains *SMC3 SMC3-AID* (BRY474), *SMC3-AID* (VG3651-3D), and *smc3-D667 SMC3-AID* (BRY482) were grown to saturation in YPD then plated in 10-fold serial dilutions onto YPD alone (YPD) or containing 0.75 mM auxin (auxin) then grown for two days at 23°C.

In the course of mapping RID mutations, we found ten RIDs within the Smc3p hinge domain (Figure 1A). Dimerization of the Smc1p and Smc3p hinges forms a toroidal structure with two interfaces termed “North” and “South” (Mishra et al. 2010). Mutations that disrupt the hinge interfaces or that neutralize the positively charged amino acids in the central channel have been studied previously (Kurze et al. 2011). Our screen identified three RIDs that mapped to the North hinge interface while six mapped near the South interface. Of the six RIDs near the South interface, five were located at or immediately adjacent to conserved glycine amino acids known to be necessary for SMC hinge dimerization *in vitro* (Figure 1B) (Hirano et al. 2001). The sixth RID, encoding an insertion of five amino acids (AAAAD) following D667, maps to a hairpin loop extending from the top of a beta-sheet that contributes to the South hinge interface. We hypothesized that the unusual position of the D667 RID might reveal a novel function of the hinge in cohesin function.

The RID screen utilizes over-expression to generate a dominant phenotype. We wanted to determine whether *smc3-D667* could support viability when expressed at native levels. For this purpose, we transformed a haploid strain bearing *SMC3-3V5-AID* as the sole *SMC3*, henceforth abbreviated *SMC3-AID*, with either an integrating *smc3-D667* or *SMC3* wild-type allele under native expression at the *LEU2* locus. We then compared growth of the *SMC3-AID* parent alone to derivatives containing either *smc3-D667* or wild-type *SMC3.* Strains were grown to stationary phase in YPD then plated as 10-fold serial dilutions on YPD media alone or containing auxin. The auxin-inducible degron (AID) epitope on Smc3-AIDp allows its rapid and specific proteasome-mediated degradation when cells are treated with auxin (Nishimura et al. 2009). As expected, the *SMC3-AID* parent is unable to grow on auxin-containing media whereas the *SMC3* wild-type containing strain shows robust growth on auxin (Figure 1C). The *smc3-D667* containing cells failed to grow on media containing auxin. The fact that *smc3-D667 SMC3-AID* cells grew well in the absence of auxin indicated that *smc3-D667* is recessive unless over-expressed. Thus, smc3-D667p was unable to support one or more essential cohesin functions.

The inviability of *smc3-D667* cells could be due to a failure of cohesin to bind DNA or a failure to perform an essential cohesin function after binding DNA. To distinguish between these possibilities, we first assessed whether smc3-D667p cohesin binds DNA. Strains containing *SMC3-AID* alone or also a second *SMC3*, either wild-type *SMC3* or *smc3-D667* were arrested in G1 phase, treated with auxin to deplete Smc3-AIDp. Cells were then synchronously released from G1 into YPD media containing auxin and nocodazole to re-arrest them in mid-M phase while maintaining Smc3-AIDp depletion (Figure 2A and Materials and Methods). To assess qualitatively whether *smc3-D667* supported binding of cohesin to chromosomes, we processed mid-M phase arrested cells for chromosome spreads and assessed chromosomal binding of the cohesin subunit Mcd1p by immunofluorescence. Mcd1p is a marker for the cohesin complex since Mcd1p cannot bind chromosomes unless it is part of the four-subunit complex (Toth et al. 1999). As expected, robust Mcd1p signal was observed on chromosome spreads from cells with Smc3p (*SMC3 SMC3-AID*) but not from cells without it (*SMC3-AID*) (Figure 2B). In *smc3-D667 SMC3-AID* cells, Mcd1p bound to chromosomes at levels similar to wild-type cells. This result indicated that smc3-D667p supports both cohesin complex assembly and binding to chromosomes.

**Figure 2:**
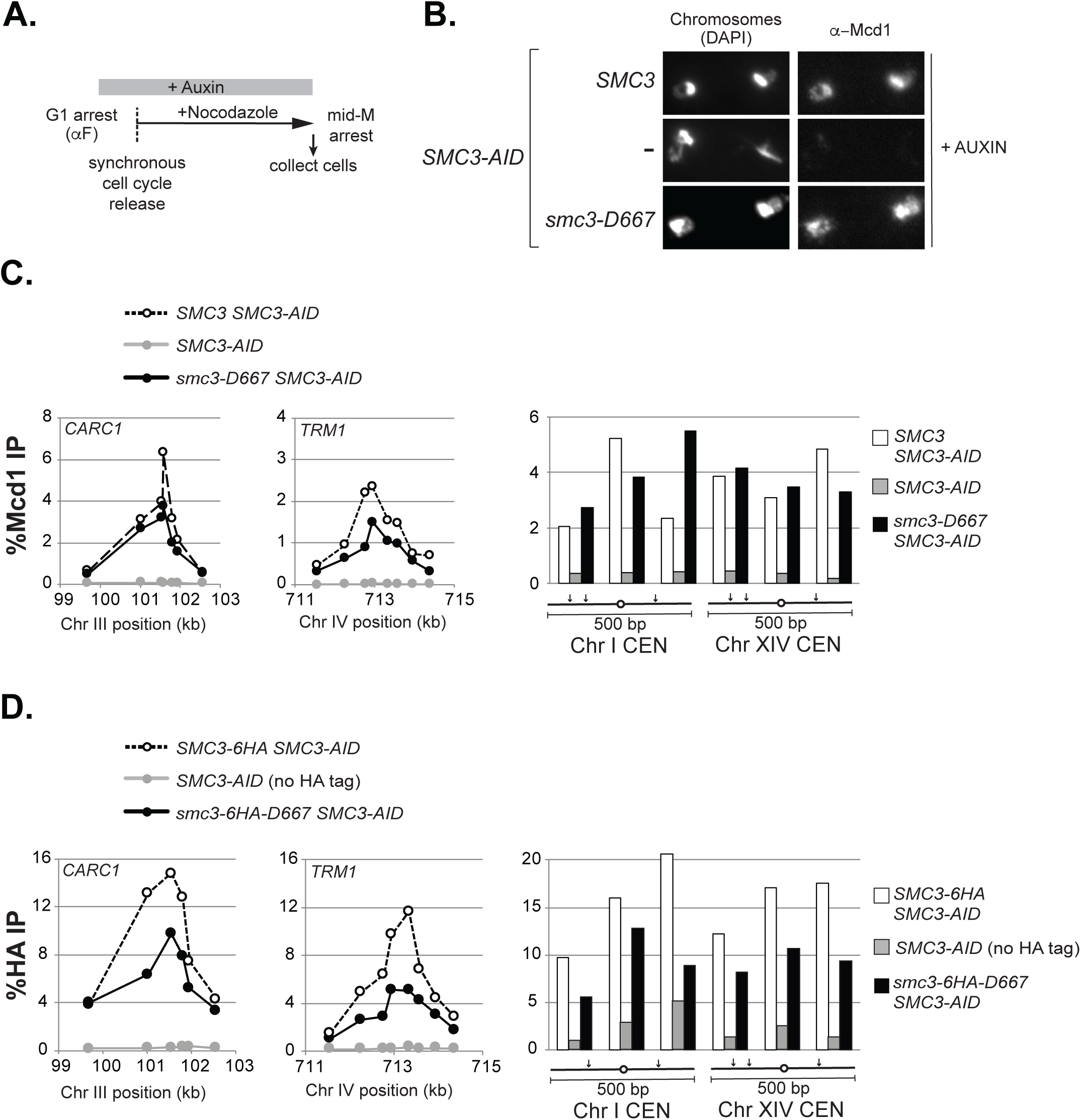
Cohesin containing smc3-D667p binds to chromosomes in mid-M phase arrested cells. A. Regimen used to prepare cells synchronously arrested in mid-M phase. Cultures were grown to mid-log phase at 23°C, treated with alpha factor for three hours to arrest cells in G1 phase then auxin was added and cells incubated an additional hour in G1 to deplete Smc3-3V5-AIDp. Cells were synchronously released from G1 arrest into YPD media containing auxin and nocodazole to re-arrest in mid-M phase (Materials and methods). B. Chromosome spreads showing that smc3-D667p cohesin binds chromosomes at levels similar to wild-type. Haploid *SMC3 SMC3-AID* (BRY474), *SMC3-AID* (VG3651-3D), and *smc3-D667 SMC3-AID* (BRY482) cells were grown as described in (A). Aliquots of mid-M phase arrested cells were fixed and processed for chromosome spreads. Bulk chromosomal DNA (DAPI) and cohesin binding (α-Mcd1) are shown. C-D. ChIP showing that *smc3-D667* cohesin binds to CARs and centromeres. C. Haploid strains in (B) were arrested in mid-M phase as described in (A) then fixed and processed for ChIP as described in materials and methods. ChIP of Mcd1p binding at *CARC1* (left) and *TRM1* (middle) and at two centromeres (right). Wild-type strain *SMC3* (dotted lines and white bars), *smc3-D667* strain (black lines and black bars) and *SMC3-AID* alone (grey lines and grey bars). (D). ChIP of HA epitope tagged Smc3p and smc3-D667p at *CARC1* (left), *TRM1* (middle) and at two centromeres (right). Haploid strains *SMC3-6HA SMC3-AID* (BRY604; dotted lines and white bars), *smc3-6HA-D667 SMC3-AID* (BRY602; black lines and black bars) and *SMC3-AID* only (VG3651-3D; grey lines and grey bars) were arrested and processed for ChIP as described in (C).

We used chromatin immunoprecipitation (ChIP) to assess whether the cohesin chromosomal binding observed via spreads reflected specific binding to CARs and centromeres. Mid-M phase cells prepared as described for chromosome spreads (Figure 2A) were fixed and processed for ChIP (Figure 2A and Materials and Methods). Cohesin binding was assessed using anti-Mcd1p antibodies. As expected, Mcd1p binding to CARs and centromeres was robust in cells with Smc3p (*SMC3 SMC3-AID*) and absent in those without it (*SMC3-AID*) (Figure 2C). Mcd1p binding in *smc3-D667* cells was similar to Wild-type at centromeres (Figure 2C, right) and at the pericentromeric *CARC1* peak (Figure 2C, left), but somewhat reduced at centromere-distal *TRM1* and *CARL1* peaks (Figure 2C, center and Supplemental Figure 2A). These results indicated that smc3-D667p cohesin localizes to CARs and centromeres. To corroborate further the DNA binding of smc3-D667p, we generated strains bearing Smc3p and smc3-D667p tagged with a 6HA epitope in the *SMC3-AID* background. Mid-M phase auxin-treated cells were prepared (Figure 2A) and the presence of smc3-6HA-D667 and Smc3-6HAp were confirmed by Western blotting (Supplemental Figure 3). We then performed ChIP using anti-HA to directly monitor the Smc3p cohesin subunit. As was observed in the Mcd1p ChIP, smc3-6HA-D667p (*smc3-6HA-D667 SMC3-AID*) bound to CARs and centromeres, albeit somewhat reduced compared to wild-type Smc3p (Figure 2D and Supplemental Figure 2B). These data, using two different cohesin subunits, show that smc3-D667p cohesin complex binds to CARs and centromeres but at reduced levels compared to wild-type.

**Figure 3:**
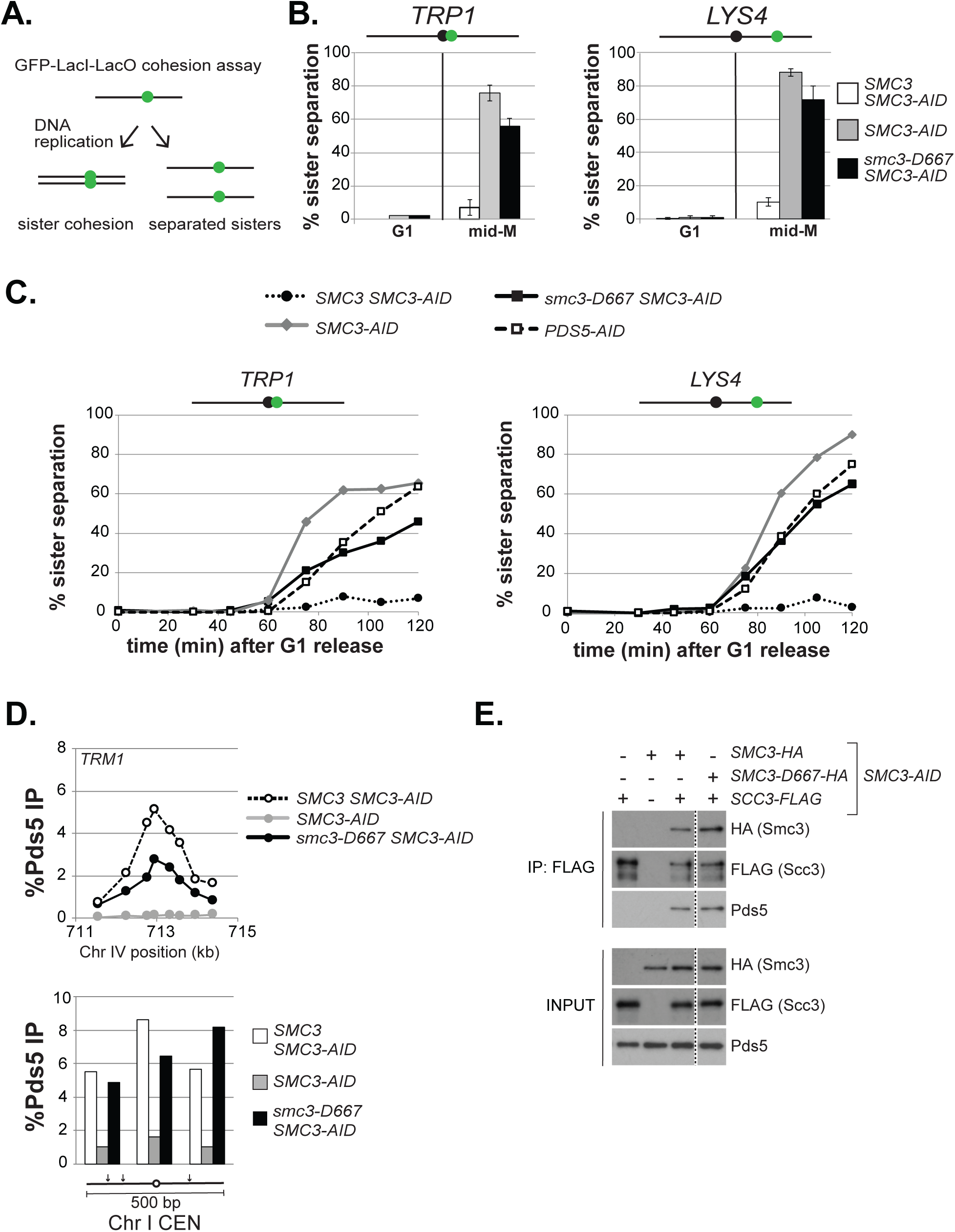
The *smc3-D667* mutant exhibits a cohesion maintenance defect. A. Schematic of cohesion loss assay using loci tagged with GFP-LacI. After replication, cells with cohesion have a single GFP focus whereas cells where cohesion is lost have 2 GFP foci. B. Cohesion loss at *CEN*-proximal *TRP1* and *CEN*-distal *LYS4* loci in mid-M phase arrested cells. Haploid strains were arrested in G1, depleted of Smc3p-AID then synchronously released from G1 and re-arrested in mid-M phase under depletion conditions as described in Figure 2A. LacO arrays integrated at *TRP1* (left) in haploid *SMC3-AID* yeast alone (BRY676) or also containing wild-type *SMC3* (BRY678), or *smc3-D667* (BRY680). LacO arrays integrated at *LYS4* (right) in *SMC3-AID* yeast alone (VG3651-3D) or containing wild-type *SMC3* (BRY474), or *smc3-D667* (BRY482). Samples were collected from G1 arrested auxin treated cells and mid-M phase arrested cells and scored for cohesion. The percentage of cells with two GFP foci (sister separation) were averaged from two independent experiments and plotted. 100-200 cells were scored per sample at each time point. Error bars represent SD. C. Time course to assess the kinetics of cohesion loss. Haploid strains were arrested in G1, treated with auxin, and synchronously released into mid-M phase arrest in auxin containing media as described in Figure 2A. Samples were collected in G1 and every fifteen minutes starting thirty minutes after G1 release and fixed to assess cohesion loss and DNA content. Data is shown as the percentage of cells with separated sisters. 100 to 200 cells were scored for cohesion for each time point. DNA content was assessed by flow cytometry and shown in Supplemental Figure 3B,C. Left side shows cohesion loss at the *CEN*-proximal *TRP1* locus. Haploid strains *SMC3 SMC3-AID* (BRY678), *SMC3-AID* (BRY676), *smc3-D667 SMC3-AID* (BRY680) and *PDS5-AID* (BRY815). Right side shows cohesion loss at the *CEN*-distal *LYS4* locus. Haploid strains *SMC3 SMC3-AID* (BRY474)*, SMC3-AID* (VG3651-3D), *smc3-D667 SMC3-AID* (BRY482) and *PDS5-AID* (TE228). D. ChIP to assess Pds5p binding to chromosomes. Haploid strains *SMC3 SMC3-AID* (BRY474), *SMC3-AID* (VG3651-3D) and *smc3-D667 SMC3-AID* (BRY482) arrested in mid-M phase according to the regimen in Figure 2A were fixed and processed for ChIP using polyclonal anti-Pds5p antibody. Pds5p binding was assessed at the CAR *TRM1* (top), and centromeres I and XIV (bottom). E. Smc3-D667p supports assembly of cohesin containing Pds5p and Scc3-3FLAGp. Haploid strains *SMC3-AID* (VG3561-3D), *SCC3-3FLAG SMC3-AID* (BRY607), *SMC3-6HA SMC3-AID* (BRY604), *SCC3-3FLAG SMC3-6HA SMC3-AID* (BRY621) and *SCC3-3FLAG smc3-6HA-D667 SMC3-AID* (BRY625) cells were grown as described in Figure 2A. Protein extracts were made and Scc3p immunoprecipitated using anti-FLAG antibody, subjected to SDS-PAGE and Western blot analysis using the indicated antibodies. Dotted line indicates where an irrelevant lane was removed.

### The D667 region of the Smc3p hinge is required to maintain cohesion

Smc3-D667p cohesin binds chromosomes, so we assayed whether it can perform cohesin’s function of tethering sister chromatids. Therefore, we assessed sister chromatid cohesion at centromere-proximal (*TRP1)* or centromere-distal (*LYS4*) loci by integrating tandem LacO repeats in strains that express a GFP-LacI fusion (Figure 3A and Materials and Methods). Strains bearing *SMC3-AID* alone or also containing either wild-type *SMC3* or *smc3-D667* were arrested in G1, treated with auxin to degrade Smc3-AIDp then synchronously released from G1 into media containing auxin and nocodazole to allow progression through S phase and arrest in mid-M phase (Figure 2A). Nearly all G1 cells in all strains contained a single GFP focus, indicating no preexisting aneuploidy (Figure 3B). As expected, only a small fraction of mid-M phase arrested cells with Smc3p (*SMC3 SMC3-AID*) lost cohesion at *TRP1* or *LYS4*, whereas cells lacking Smc3p (*SMC3-AID*) had almost complete loss of cohesion. Nearly two-thirds of cells expressing only *smc3-D667* (*smc3-D667 SMC3-AID)* also had lost cohesion at these two loci. This result suggested that the D667 region of the hinge was required for either robust establishment and/or maintenance of cohesion.

These two possibilities can be distinguished by kinetic analysis of cohesion in populations of cells synchronously progressing through the cell cycle. Mutants that compromise cohesion establishment like those defective in core subunits of cohesin *MCD1*, *SMC3,* and *SMC1* exhibit sister chromatid separation immediately after DNA replication (Guacci et al. 1997; Michaelis et al. 1997). Mutants that compromise cohesion maintenance like those defective in the cohesin regulator *PDS5* also lose cohesion but significantly later in the cell cycle than establishment mutants (Tanaka et al. 2001; Stead et al. 2003; Noble et al. 2006; Eng et al. 2014). Using the same strains as described above along with a *PDS5-AID* strain, we assessed when cohesion was lost in *smc3-D667*. Strains were arrested in G1 and treated with auxin to degrade Smc3-AIDp, then released from G1 in the presence of auxin and nocodazole to allow cells to progress through S phase and arrest in mid-M. After release from G1, aliquots of cells were removed every fifteen minutes to assess DNA content and cohesion at *TRP1* and *LYS4* (Figure 3C).

From analysis of the DNA content, all strains exhibited nearly identical kinetics of progression through S phase and subsequent arrest in mid-M (Supplemental Figure 4A). As expected for cells expressing Smc3p (*SMC3 SMC3-AID*), sister chromatids were paired through mid-M arrest so few cells with separated sisters were detected. In contrast, both strains lacking Smc3p (*SMC3-AID*) and Pds5p (*PDS5-*AID) lost cohesion. However, the cohesion loss in the *PDS5-AID* cells was delayed by about 20 minutes, as published previously (Eng et al. 2014). Cells expressing only smc3-D667p (*smc3-D667 SMC3-AID*) resembled *PDS5-AID* cells, with delayed cohesion loss at the *LYS4* locus and a more pronounced delay in cohesion loss at the *TRP1* locus. This delay in cohesion loss in cells with smc3-D667p demonstrated that *smc3-D667* cells, like Pds5p-deficient cells, could establish but not maintain cohesion. Thus, the D667 region of the Smc3p hinge is important specifically for efficient maintenance of cohesion at both *CEN*-proximal and *CEN*-distal loci.

**Figure 4:**
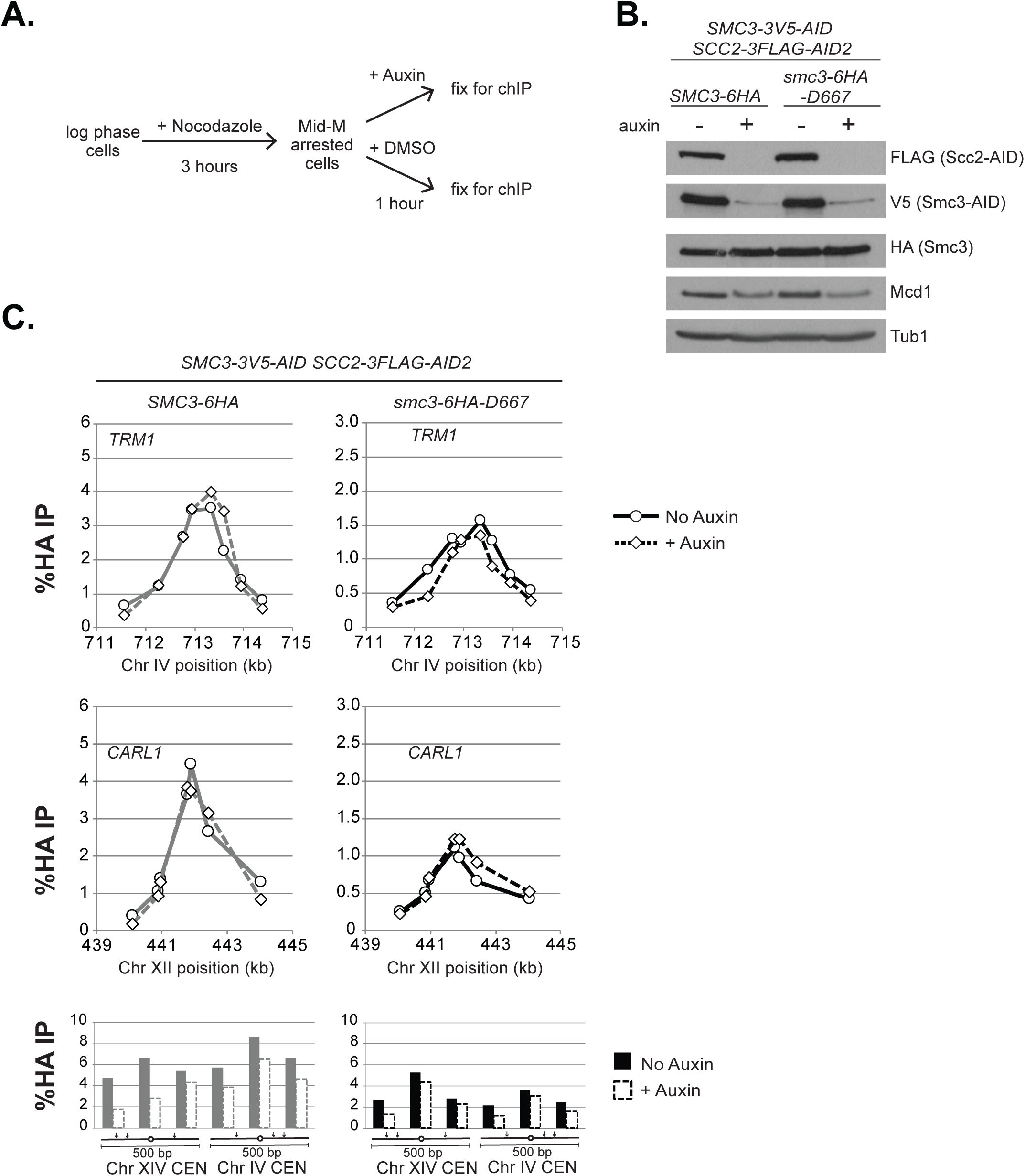
*smc3-D667* supports stable cohesin binding to chromosomes A. Regimen used to assess stability of cohesin binding to DNA upon depletion of the loader subunit Scc2p. Haploid *SMC3-3V5-AID SCC2-3FLAG-AID2* strains expressing either *SMC3-6HA* (BRY839) or *smc3-6HA-D667* (BRY841) were grown to mid-log phase and arrested in mid-M phase by incubation with nocodazole for three hours. Cultures were split and auxin added to one half then both halves incubated for one hour. Cells aliquots were collected to make protein extracts or fixed and processed for ChIP (Materials and Methods). B. Western Blot analysis showing depletion of AID tagged proteins. Protein extracts (TCA lysed) of strains in (A) were subjected to SDS-PAGE and analyzed by Western blot. Depletion of Scc2p-3FLAG-AID (FLAG) and Smc3p-3V5-AID (V5) is shown. Antibodies assessing levels of Smc3p (HA) and Mcd1p (Mcd1) cohesin subunits and a loading control (Tub1). C. ChIP to assess the stability of cohesin (Smc3p) binding at CARs and centromeres. Cultures of strains from (A) were fixed and processed for ChIP. Smc3-6HAp binding (left side) and smc3-6HA-D667p binding (right side) at CARs and centromeres in control cells (solid lines and filled columns) and auxin-treated cells depleted for Scc2-3FLAG-AID2p and Smc3-3V5-AIDp (dashed lines and open columns). From top to bottom: binding to CARs *TRP1* and *CARL1*, and centromeres XIV and IV.

Cohesin is required to recruit the maintenance factor Pds5p to chromosomes (Hartman et al. 2000; Panizza et al. 2000). Since cells expressing smc3-D667p displayed a cohesion maintenance defect identical to cells depleted of Pds5p, we tested whether smc3-D667p cohesin was able to recruit Pds5p to chromosomes. To address this possibility, we first analyzed whether smc3-D667p supported Pds5p binding to chromosomes by ChIP using a Pds5p antibody (Figure 3D and Supplemental Figure 4B). The ratio of Pds5p bound to CARs and centromeres in cells with smc3-D667p (*smc3-D667 SMC3-AID*) to Smc3p was very similar to that seen for Mcd1p or smc3-6HA-D667p. These results indicate that cohesin with smc3-D667p can bind Pds5p and recruit it to chromosomes. The ability of Pds5p to bind cohesin with smc3-D667p was then tested by co-immunoprecipitation (Figure 3E). Cells expressing FLAG-tagged Scc3p and HA-tagged Smc3p or smc3-D667p were arrested in M-phase after auxin-mediated depletion of Smc3-AIDp. Scc3p was immunoprecipitated using anti-FLAG antibody and cohesin subunits detected in the precipitates by Western blot. As expected, no Pds5p was detected in the FLAG immunoprecipitate from cells lacking Smc3p or when Scc3p was untagged (first and second lanes), while Pds5p and smc3-6HAp were detected in the immunoprecipitate from cells expressing Smc3-6HAp (third lane). Importantly, similar Pds5p levels were observed in the immunoprecipitate from cells expressing smc3-D667-6HAp (fourth lane). Thus, smc3-D667p cohesin binds Pds5p and recruits it to DNA.

### The D667 region of the Smc3p hinge is not required for its stable binding to chromosomes

Cohesin is known to convert from a DNA-bound, untethered state to a tethered state in S phase (Ünal et al. 2008; Ben-Shahar et al. 2008). We envisioned two models by which cohesion that had been established in S phase by smc3-D667p could fail to be maintained as cells progressed into M phase. In one model, cohesin reverts back to its untethered state without perturbing cohesin binding to DNA. Precedence for this phenotype comes from the cohesin mutant *mcd1-ROCC* which, like *smc3-D667*, is defective for cohesion maintenance (Eng et al. 2014). Alternatively, the smc3-D667p is less stably bound so dissociates from DNA. In this model, following cohesion establishment, cohesin dissociation from chromosomes could manifest as a cohesion maintenance defect. Detecting putative cohesin dissociation is difficult, because the Scc2p/Scc4p complex continues loading cohesin onto chromosomes in mid-M phase creating a pool of bound cohesin that does not contribute to cohesion (Lengronne et al. 2006). Therefore, the Scc2p/Scc4p complex must be inactivated to allow detection of cohesin dissociation.

To distinguish between these two models, we examined the stability of smc3-6HA-D667p binding to DNA under conditions where additional loading was prevented by depletion of the cohesin loader subunit Scc2p. This loader depletion approach revealed that in wild-type cells, cohesin (Mcd1p) binds stably at CARs but exhibits reduced stability at centromeres (Eng et al. 2014). Therefore, we replaced *SCC2* with *SCC2-3FLAG-AID* in *SMC3-AID* strains bearing either wild-type Smc3-6HAp or smc3-D667-6HAp. Cultures of these strains were grown to mid-log phase and arrested in mid-M phase by addition of nocodazole. Cultures were then split and either auxin or vehicle (DMSO) added, then incubated for one hour. The aliquot containing auxin will deplete both Scc2-3FLAG-AIDp and Smc3-3V5-AIDp. Samples were collected and either fixed for ChIP or processed for Western Blot analysis (Figure 4A). Depletion of Scc2-3FLAG-AIDp and Smc3-3V5-AIDp was confirmed by Western blot (Figure 4B).

ChIP of Smc3-6HAp showed no difference in binding to CAR peaks *TRM1* and *CARL1* after Scc2-3FLAG-AIDp depletion (Figure 4C, left). The persistence of high ChIP levels even after an hour indicated that cohesin remained very stably bound to DNA. Similarly, smc3-6HA-D667p ChIP at *TRM1* and *CARL1* peaks was unchanged by Scc2-3FLAG-AIDp depletion (Figure 4C, right). At centromeres, Smc3-6HAp shows somewhat reduced binding after Scc2-3FLAG-AIDp depletion, confirming this cohesin is less stably bound. Similarly, somewhat reduced binding of smc3-6HA-D667p to centromeres was observed. These results demonstrated that smc3-6HA-D667p was as stably bound to chromosomes as wild-type Smc3-6HAp. Importantly, our results indicated that in mid-M phase arrested *smc3-D667* cells, when most sister chromatid cohesion is lost (Figure 3), smc3-D667p cohesin is stably bound to chromosomes. Thus, the D667 region of the Smc3p hinge performs a function in maintaining cohesion other than ensuring stable binding to DNA.

### The D667 region of the Smc3p hinge modulates cohesion and supports viability by a mechanism independent of Eco1p-dependent acetylation

Eco1p is necessary for establishing cohesion during S phase through its acetylation of Smc3p at lysines K112 and K113. Although cohesion establishment occurs during S phase, Smc3p acetylation remains until anaphase onset, suggesting it may be required to maintain cohesion (Beckouet et al. 2010). Since smc3-D667p supported cohesion establishment, we predicted that it would be acetylated by Eco1p. Therefore, we used an antibody that specifically recognizes acetylated Smc3p-K113 to test the acetylation of smc3-D667p in cells arrested in mid-M. Cells were arrested in mid-M after auxin depletion (Figure 5A). As expected, in cells depleted of Eco1-AIDp or Smc3-AIDp, no acetylated Smc3p was detected (Figure 5B). While wild-type Smc3p showed strong acetylation signal, acetylation signal for smc3-D667p was reduced. A reduction in acetylation signal was expected because cohesin was known to be acetylated only after binding to DNA and less cohesin with smc3-D667p was bound to DNA than wild-type cohesin (Figure 2). Direct comparison of acetylation levels is possible when signal from the acetylation-recognizing antibody is linear across the observed range. However, we found that signal from the acetylation antibody was non-linear (Supplementary Figure 5), making it possible that smc3-D667p acetylation levels were closer to Smc3p than Figure 5B suggested.

**Figure 5:**
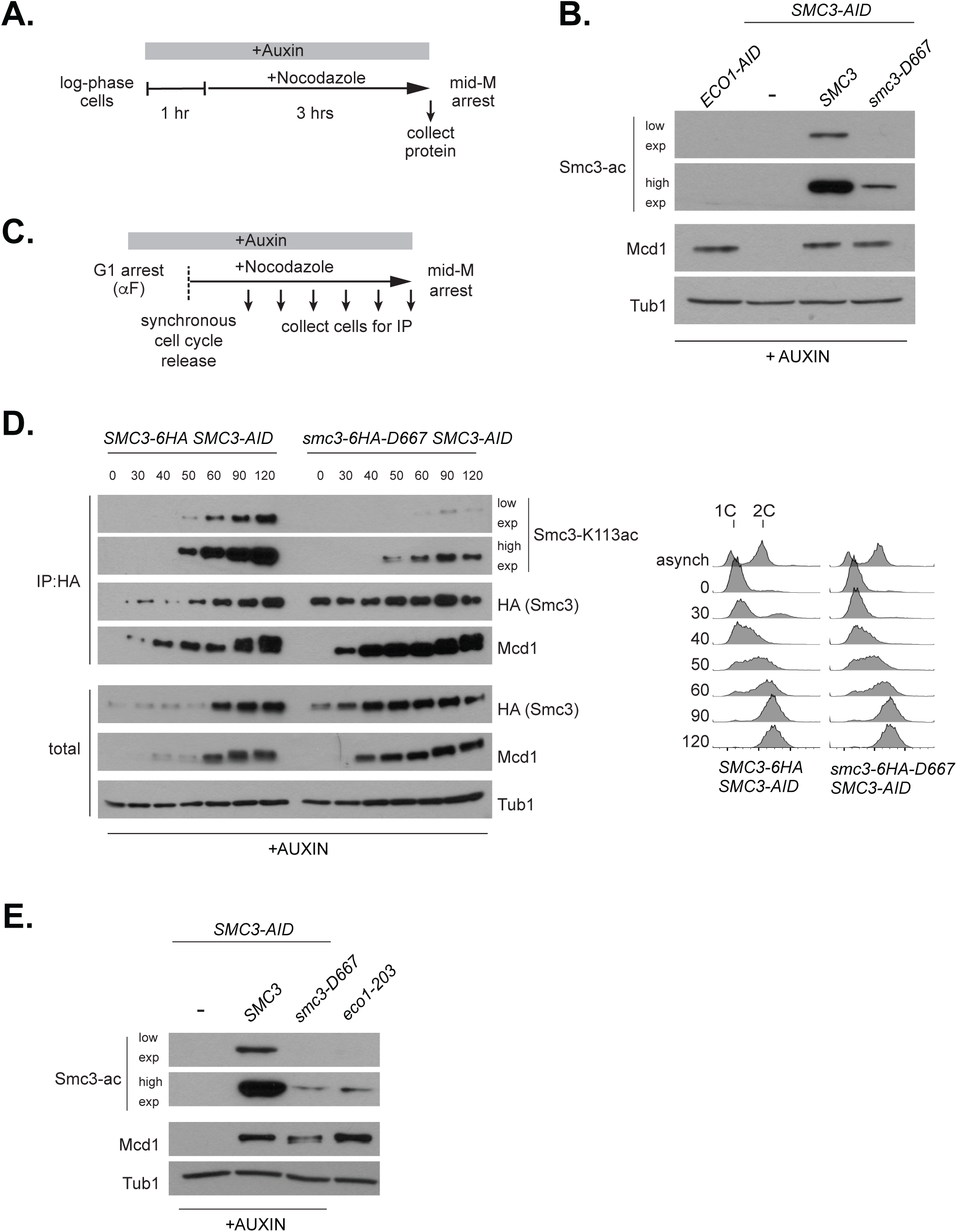
smc3-D667p has reduced acetylation at K113 A. Regimen used to assess Smc3-K113 acetylation in mid-M phase arrested cells. Early log phase cultures were treated with 0.75 mM auxin for one hour to deplete Smc3-3V5-AIDp then nocodazole was added and cultures incubated three hours to arrest cells in mid-M phase. B. Reduced K113 acetylation of smc3-D667p. Haploid *ECO1-AID* (VG3633-2D), *SMC3-AID* (VG3651-3D), *SMC3 SMC3-AID* (BRY474), and *smc3-D667 SMC3-AID* (BRY482) cultures grown as described in (A). Protein extracts were made and subjected to SDS-PAGE then analyzed by Western blot. Antibodies against Smc3-K113 acetylation (Smc3-ac) are shown as short and long exposures, anti-Mcd1p antibodies (Mcd1p) serve as control for cohesin levels and antibodies against tubulin (Tub1) for a loading control. C. Regimen used to determine the kinetics of Smc3-K113 acetylation establishment within a single cell cycle. Log phase cultures grown in YPD at 23°C were arrested in G1 using alpha factor, treated with auxin to deplete Smc3p-AID in G1 then released into fresh YPD containing auxin and nocodazole to synchronously arrest cells in mid-M phase (Materials and methods). D. smc3-6HA-D667p has reduced acetylation in S phase but acetylation remains in mid-M phase. Haploid *SMC3-AID* cells expressing Smc3-6HAp (BRY604, left) or smc3-6HA-D667p (BRY602, right) were grown as described in (C). Aliquots were taken at the indicated time points and protein extracts made. A small portion was reserved for total protein then anti-HA antibody added to immunoprecipitate Smc3-6HAp or smc3-6HA-D667p (Materials and Methods). Samples were subjected to SDS-PAGE then analyzed by Western blot. Antibodies against Smc3-K113 acetylation (Smc3-ac) and both short and long exposure shown for better comparison. Antibodies were used to monitor levels of the Smc3p (HA) and Mcd1p (Mcd1) cohesin subunits and anti-Tubulin antibodies (Tub1) used as a loading control. Samples were also collected to assess DNA content by flow cytometry (right side). E. Similar levels of K113 acetylation in *smc3-D667* and *eco1-203* at permissive temperature. Haploid strains *SMC3-AID* (VG3651-3D), *SMC3 SMC3-AID* (BRY474), *smc3-D667 SMC3-AID* (BRY482) and *eco1-203* (VG3506-5D) were treated as described in (A). Protein extracts were made, subjected to SDS-PAGE and Western blot analysis. Antibodies against Smc3-K113 acetylation (Smc3-ac) and both short and long exposure shown for better comparison. Anti-MCD1 antibodies (Mcd1) were used as a control for cohesin levels and anti-Tubulin antibodies (Tub1) for a loading control.

To assess whether the reduced amount of smc3-D667p acetylation was responsible for the cohesion maintenance defect, we first asked whether a change in acetylation levels correlated with the appearance of the cohesion defect. Reduced smc3-D667p acetylation may have resulted from a failure to acetylate it in S phase or to maintain it after S phase. To distinguish between these possibilities, we immunoprecipitated smc3-6HA-D667p from cells progressing synchronously through S phase following release from G1 arrest (Figure 5C). As expected, wild-type Smc3-6HAp acetylation began to appear during S phase then increased and remained high through M phase arrest (Figure 5D). While acetylation of smc3-6HA-D667p was lower than WT in early S phase, it increased as cells progressed into M phase. Therefore, *smc3-D667* cells establish cohesion with low smc3-D667p acetylation levels but its failure to maintain cohesion is not due to a subsequent decrease in acetylation levels.

We further examined the correlation between Smc3p acetylation levels and cohesin function by asking whether low levels of Smc3p acetylation always led to loss of essential cohesin function. Temperature-sensitive *eco1* mutants (*eco1-203* and *eco1-1*) establish and maintain cohesion at permissive temperature yet *eco1-1* has greatly reduced acetylation (Toth et al. 1999; Rowland et al. 2009; Heidinger-Pauli et al. 2009). We therefore compared Smc3p acetylation levels of the *eco1-203* mutant grown at the permissive temperature 23°C to the *smc3-D667* mutant. The level of Smc3p acetylation in *eco1-203* cells was very similar to *smc3-D667* cells (Figure 5E). This result suggested that the level of smc3-D667p acetylation was sufficient to support cohesion function. However, we could not rule out that the acetylation level of smc3-D667p was below a critical threshold too subtle to distinguish by Western blot.

We sought additional support for the idea that the lower smc3-D667p acetylation level is not responsible for its mutant phenotype. For this purpose, we assayed the *smc3-D667* mutant in the *SMC1-D1164E* mutant background, as this *SMC1* allele completely bypasses the need for Smc3p acetylation in both cohesion and viability (Çamdere et al. 2015; Elbatsh et al. 2016). In the presence of auxin, *smc3-D667 SMC3-AID* and *SMC1-D1164E smc3-D667 SMC3-AID* cells were inviable (Figure 6A). Therefore, the viability defect of *smc3-D667* is distinct from *eco1-ts* and deletion mutants, which are bypassed by *SMC1-D1164E*. We next asked whether *SMC1-D1164E* restored cohesion to *smc3-D667* cells as was observed for the *eco1*Δ *wpl1*Δ mutant and *eco1*Δ cells (Çamdere et al. 2015). As expected, *SMC1-D1164E* restored cohesion at the *LYS4* locus in the *eco1*Δ *wpl1*Δ mutant (Figure 6B and Çamdere et al. 2015). However, *SMC1-D1164E* failed to restore cohesion to the *smc3-D667 SMC3-AID* mutant in the presence of auxin (Figure 6C). These results supported the idea that the viability and cohesion defects of *smc3-D667* cells were independent of reduced levels of Smc3p acetylation.

**Figure 6:**
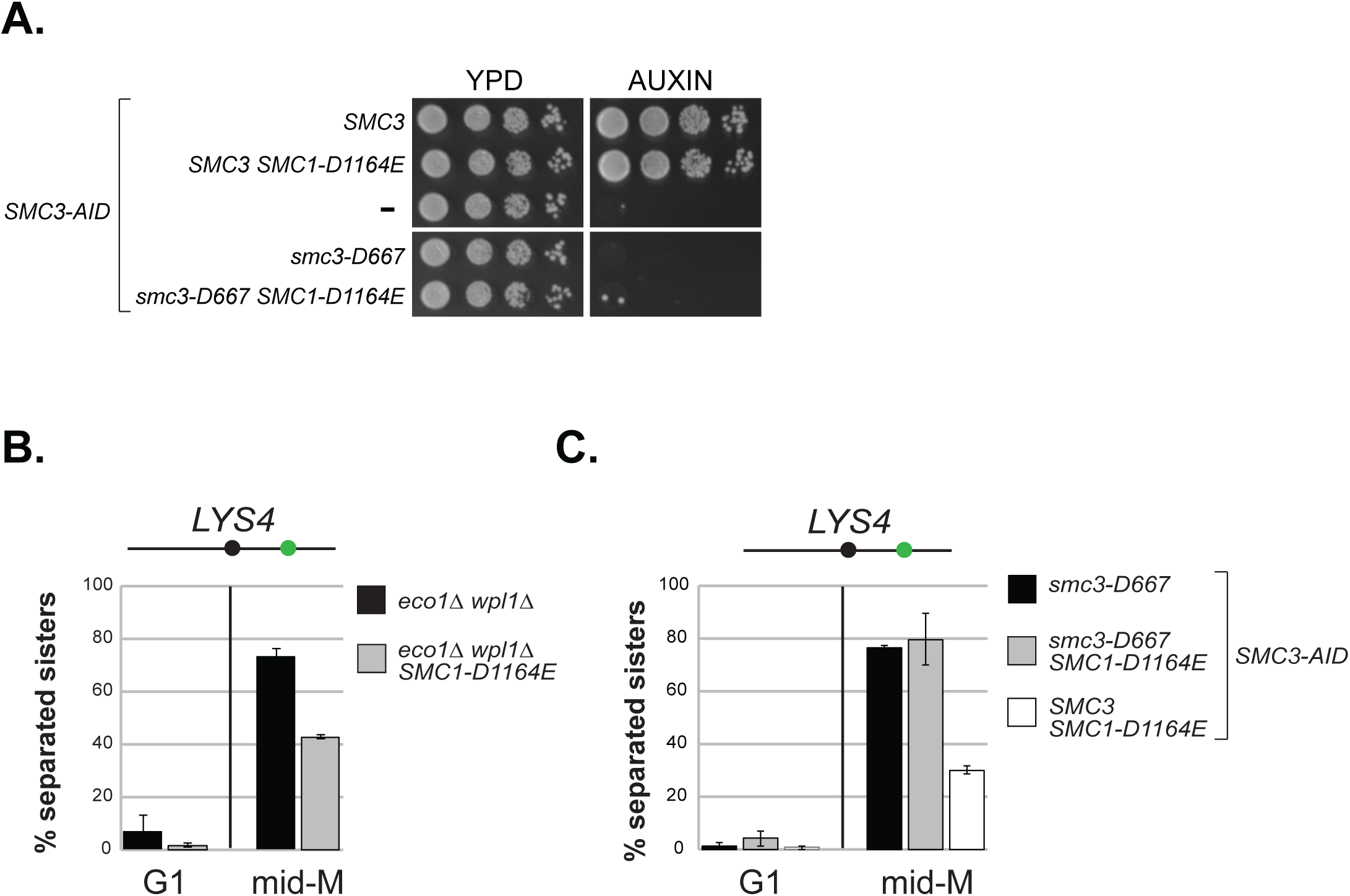
The *SMC1-D1164E* mutation fails to suppress the inviability or cohesion defect of *smc3-D667* A. *smc1-D1164E* failed to restore viability to *smc3-D667* cells. Haploid strains *SMC3 SMC3-AID* (BRY474), *SMC3 SMC3-AID SMC1-D1164E* (BRY832), *SMC3-AID* (VG3651-3D), *smc3-D667 SMC3-AID* (BRY482), and *smc3-D667 SMC3-AID SMC1-D1164E* (BRY833) were grown to saturation in YPD, then plated as ten-fold serial dilutions onto YPD alone (YPD) or containing 0.75 mM auxin (YPD + auxin) and incubated 2 days at 23°C. B. *SMC1-D1164E* suppresses cohesion loss of the *eco1*Δ *wpl1*Δ mutant in mid-M phase arrested cells. Haploid strains *eco1*Δ *wpl1*Δ (VG3503-4A), *SMC1-D1164E eco1*Δ *wpl1*Δ (VG3575-2C) grown as described in Figure 2A. Cells from G1 and mid-M phase arrest were fixed and processed and scored for cohesion loss at the *CEN*-distal *LYS4* locus. C. *SMC1-D1164E* fails to suppress cohesion loss of *smc3-D667* cells. Haploid strains *smc3-D667 SMC3-AID* (BRY482), *smc3-D667 SMC3-AID SMC1-D1164E* (BRY833), and *SMC3 SMC3-AID SMC1-D1164E* (BRY832) cells were grown according the regimen in Figure 2A and processed to assess cohesion loss at the *CEN*-distal *LYS4* locus as described in (B). For both (B) and (C), the percentage of cells with two GFP foci (sister separation) were derived from two independent experiments. 100-200 cells were scored per sample at each time point. Error bars represent SD.

### The D667 region of the Smc3p hinge is required for *rDNA* condensation and viability even in the absence of antagonism by Wpl1p

In addition to sister chromatid cohesion, cohesin and its regulators Pds5p and Eco1p are required for the proper mitotic condensation of chromatids in budding yeast (Guacci et al. 1997; Hartman et al. 2000; Skibbens et al. 1999). We addressed whether *smc3-D667* cells supported condensation by examining the morphology of the *rDNA* locus on chromosome XII. In chromosome spreads the *rDNA* is located on the periphery of the primary chromosome mass. In interphase, the *rDNA* can be seen as a diffuse puff while in M phase it condenses into a loop (Guacci et al. 1994). Chromosome spreads of the *SMC3-AID* and *PDS5-AID* strains were prepared from cells arrested in mid-M phase (Figure 7A). The *rDNA* morphology was scored as either 1) tight, fully-condensed loop 2) wide, decondensed loop or 3) diffuse, with no apparent loop. In cells with wild-type Smc3p, the *rDNA* formed tight loops in almost all chromosome masses, indicative of chromosome condensation. In cells lacking Smc3p (*SMC3-AID)*, the *rDNA* was almost always present as a diffuse mass, recapitulating the established role of Smc3p and cohesin in condensation. Cells expressing only smc3-D667p or depleted of Pds5p (*PDS5-AID*) exhibited very similar condensation defects and tight loops were rarely observed (Figure 7A). Thus, the D667 region of the Smc3p hinge is needed for two M phase functions of cohesin, the maintenance of cohesion and condensation.

**Figure 7:**
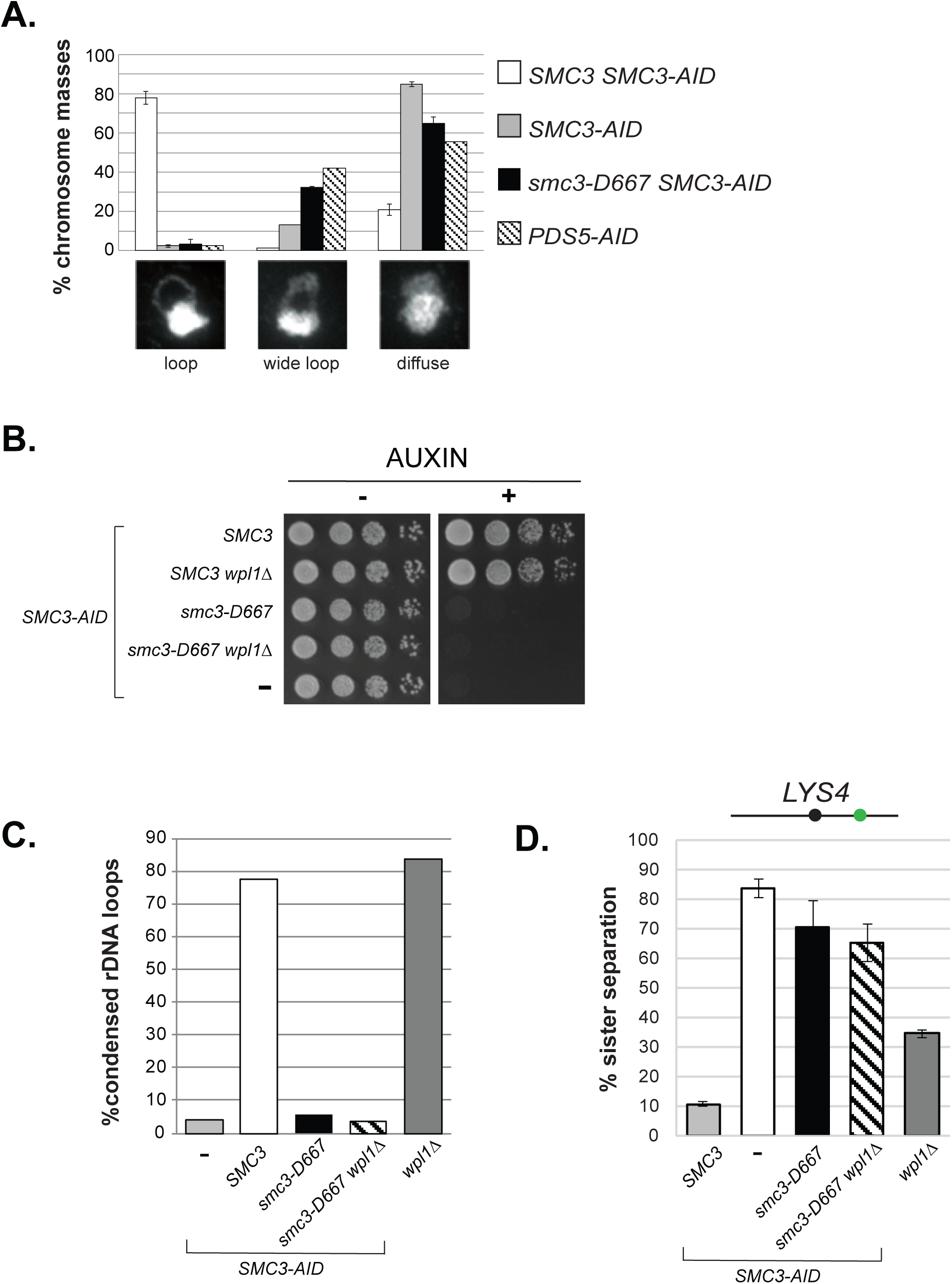
The *smc3-D667* mutant is defective in condensation and cohesion even in the absence of cohesin antagonist Wpl1p. A. Condensation of the *rDNA* locus in *smc3-D667* cells. Percentage of chromosome masses displaying tight loop, wide loop, or diffuse *rDNA* morphologies. Haploid strains *SMC3 SMC3-AID* (BRY474), *SMC3-AID* (VG3651-3D), *smc3-D667 SMC3-AID* (BRY482), and *PDS5-AID* (TE228) were grown and treated as in Figure 2A then processed as if for *in situ* hybridization (see Materials and Methods). Chromosome masses were scored for *rDNA* locus morphology after staining with DAPI. Shown are averages from two independent experiments in which 100 chromosome masses were scored. Error bars depict SD. B. *wpl1*Δ fails to restore viability to *smc3-D667* cells. Haploid *SMC3-AID* strain derivatives with *SMC3* (BRY474), *SMC3 wpl1*Δ (BRY716), *smc3-D667* (BRY482), *smc3-D667 wpl1*Δ (BRY718), or *SMC3-AID* alone (VG3651-3D) were grown and plated as described in Figure 1C. C. Quantification of condensed *rDNA* masses from mid-M phase arrested cells. Haploid strains *SMC3-AID* (VG3651-3D), *SMC3 SMC3-AID* (BRY474), *smc3-D667 SMC3-AID* (BRY482), *smc3-D667 SMC3-AID wpl1*Δ (BRY718), and *wpl1*Δ (DK5561) were treated and processed as in (A). The percentage of chromosome masses displaying a tight *rDNA* loop is shown. D. Cohesion loss in *smc3-D667 wpl1*Δ cells. Haploid *wpl1*Δ (DK5561) and *SMC3-AID* strain derivatives with *SMC3* (BRY474), *SMC3-AID* alone (VG3651-3D), *smc3-D667* (BRY482), *smc3-D667 wpl1*Δ (BRY718) were treated as in Figure 2A and the percentage of separated sisters at the *LYS4* locus plotted. Error bars represent the SD.

We next asked whether the condensation defect and inviability of *smc3-D667* cells was due to antagonism by Wpl1p. Deletion of *WPL1* restores viability to *eco1* temperature-sensitive or *eco1*Δ strains which have impaired or absent acetylation (Rowland et al. 2009; Guacci and Koshland 2012). If the defect of *smc3-D667* can be attributed to a loss of Eco1p function, then *wpl1*Δ would restore condensation and viability to *smc3-D667* cells. To test this idea, we characterized the consequences of *WPL1* deletion in the *smc3-D667* strain. *wpl1*Δ failed to restore viability to *smc3-D667 SMC3-AID* cells on media containing auxin (Figure 7B). Consistent with *smc3-D667* representing a defect distinct from cells lacking Smc3p acetylation, *wpl1*Δ failed to restore condensation of the *rDNA* or cohesion to *smc3-D667* cells (Figure 7C and 7D, respectively). Altogether, our observations confirmed that the critical defects in *smc3-D667* cells were independent of Smc3p acetylation or antagonism by Wpl1p.

### The D667 region is necessary for interallelic complementation

Interallelic complementation between alleles of *SMC3* or *MCD1* revealed the ability of two separate cohesin complexes to share activities to restore cohesin functions. Additional evidence suggests that this communication between cohesins might reflect direct cohesin-cohesin interaction on chromosomes (Eng et al. 2015). We wondered whether the D667 region of the hinge was needed for cohesin-cohesin communication. To test this idea, we asked whether *smc3-D667* could partner with the temperature sensitive *smc3-42* allele to exhibit interallelic complementation. The temperature sensitive *smc3-42* strain cannot grow at its restrictive temperature of 34°C. Previously it had been shown that the *smc3-K113R* allele cannot support viability as the sole copy of *SMC3*. However, a strain in which both *smc3-K113R* and *smc3-42* alleles are present exhibits robust growth at 34°C, a condition in which neither single mutant can grow (a summary of complementation relationships is provided in Figure 8B). With this knowledge, we asked whether *smc3-D667* could substitute for *smc3-K113R* and complement *smc3-42*. As a metric for the extent of interallelic complementation, we repeated the previous experiment with *smc3-42* and *smc3-K113*. As expected, at 34°C neither *smc3-42* nor *smc3-K113R* single mutants were viable, while the *smc3-42 smc3-K113R* double mutant showed robust growth similar to wild-type (Figure 8A). As expected, the *smc3-D667* single mutant failed to grow. The double *smc3-42 smc3-D667* mutant resembled the growth of *smc3-42* alone. Thus, the property of interallelic complementation observed between *smc3-42* and *smc3-K113R* was not observed between *smc3-42* and *smc3-D667*. Therefore, *smc3-D667* lacks the activity necessary for interallelic complementation. This result suggested that the D667 region of the hinge is necessary for cohesin-cohesin communication.

**Figure 8:**
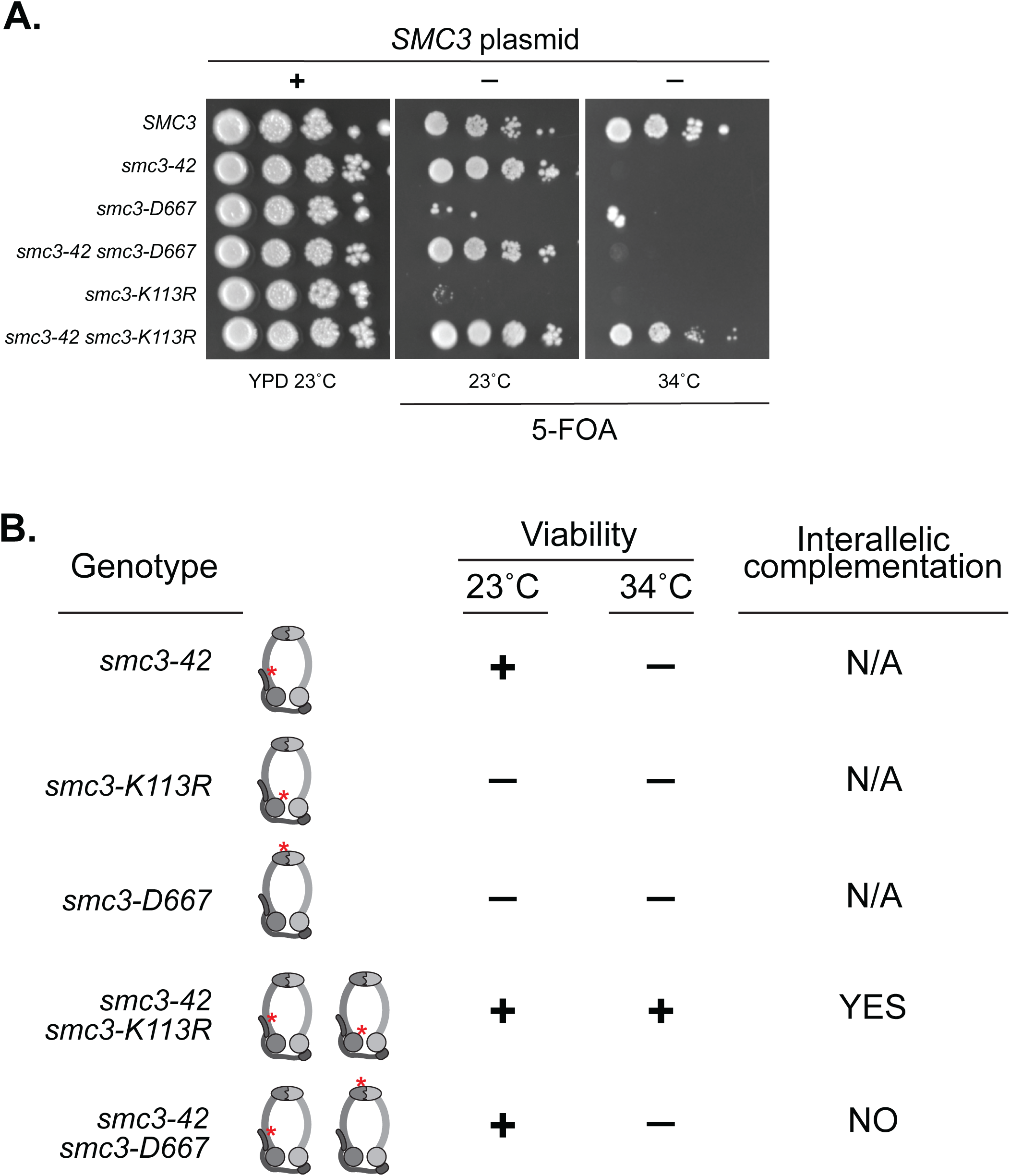
The D667 region is necessary for interallelic complementation. A. Assessing whether *smc3-D667* complements the *smc3-42* mutant. Haploid strains *SMC3* (VG3486), *smc3-42* (TE576), *smc3-D667* (BRY467), *smc3-42 smc3-D667* (BRY756), *smc3-K113R* (VG3486-K113R), and *smc3-42 smc3-K113R* (TE578) all contain the *SMC3 URA3 CEN* plasmid. Strains were grown to saturation in YPD cultures to allow loss of the *SMC3 URA3 CEN* plasmid then played at 10-fold serial dilutions on YPD or 5-FOA plates and incubated at the indicated temperatures. B. Table summarizes interallelic complementation of haploid cells harboring the temperature-sensitive *smc3-42* allele (Eng et al. 2015) and (A).

## Discussion

Cohesin has a complex architecture with a heterodimeric ATPase domain and a hinge domain connected by a long coiled coil. The roles of these domains in cohesin’s activity on chromosomes is poorly understood. Here, we identified and characterized *smc3-D667*, a mutant in the Smc3p hinge domain which blocks cohesin function in M phase. Kinetic analyses of cohesion during the cell cycle reveal that this mutation allows cohesion establishment but impairs subsequent maintenance of cohesion. We also show that this mutation impairs mitotic chromosome condensation of the *rDNA*. However, this mutation does not perturb the stable association of cohesin with chromosomes as measured by the persistence of this association even after loader inactivation. Together, our results support a function of cohesin’s hinge domain in cohesion maintenance and condensation independent of cohesin’s stable binding to chromosomes.

The cohesion maintenance and condensation functions of the hinge domain revealed by *smc3-D667* have not been reported previously. Two mutations that impact the North and South interfaces of the hinge dimer revealed a role of the hinge in cohesin binding to chromosomes, as expected given the role of the hinge dimer in maintaining the topological integrity of cohesin (Mishra et al. 2010). The novel phenotypes of *smc3-D667* are consistent with D667 localization, determined by alignment to Smc3p homologs, within a loop not expected to impact the dimer interface. One study designed a cluster of mutations in *SMC1* and *SMC3* that neutralize the positive charges in a central channel formed by hinge dimerization (Kurze et al. 2011). This cluster of mutations (charge neutralization alleles) caused defects in cohesion and Smc3p acetylation but did not impair stable binding of cohesin to chromosomes, all phenotypes similar to the *smc3*-*D667* allele. However, unlike our study of *smc3-D667,* the charge neutralization alleles were not analyzed for establishment and maintenance cohesion, the functional significance of the reduced Smc3p acetylation, or condensation. If these alleles had the same cohesion and condensation defects as the *smc3-D667* allele, as we predict, these results would imply that changes to two distinct regions of the hinge dimer contribute to a common function needed for cohesion maintenance and condensation. The potential cooperation of the D667 region of the Smc3p hinge and the hinge channel could reflect a previously unrecognized conformational change of the hinge dimer needed for cohesin function. Indeed, in addition to the strict toroidal structures seen by crystallization of the cohesin or *Tm*SMC hinge dimers, a recently published structure of the related *Gs*SMC hinge dimer revealed that hinge dimers may adopt an asymmetric, relaxed conformation resembling a spring washer (Haering et al. 2002; Kurze et al. 2011; Kamada et al. 2017). Surprisingly, while both hinge interfaces remained intact in this structure, the relaxed face of the GsSMC hinge dimer involved a break in the beta sheet connected by a loop homologous to the D667 loop of Smc3p. Together with our results, further investigation of hinge structural flexibility on conformations and functions of cohesin seem worthwhile.

The unusual phenotypes of *smc3-D667* are also strikingly similar to those described for Pds5p depletion and *mcd1* alleles (Chan et al. 2013, Eng et al. 2014). They all allow stable cohesin binding to DNA but cause defects in cohesion maintenance and condensation. These common phenotypes suggest that the hinge, Pds5p and Mcd1p cooperate in a common molecular function. Indeed, this common function provides a biological explanation for in vivo FRET studies that suggest the formation of a complex involving the head, hinge, and Pds5p (Mc Intyre et al. 2007), and recent biochemical experiments that detected a supramolecular complex between the *S. pombe* hinge dimer and Psc3p (Scc3p ortholog) which binds to the head-associated Rad21p (Mcd1p ortholog) and Pds5p. Altogether these biochemical results along with our study support the idea that the hinge, Mcd1p, and Pds5p cooperate in a structural conformation required to promote cohesion maintenance and condensation.

Potential insight into the molecular function of this complex conformation comes from several additional observations. One possibility was the protection of Eco1p acetylation of Smc3p. Here, we show that while the level of smc3-D667 acetylation is lower than wild-type, it is equal to that of the *eco1-203* mutant at its permissive temperature, which supports both viability and sister chromatid cohesion. Furthermore, we show that *SMC1-D1164E* and *wp1*Δ, two different mutations previously shown to bypass the absence of Eco1p acetylation in viability, cohesion (only *smc1-D1164E*) and condensation (only *wpl1*Δ) are unable to restore these functions to the *smc3-D667* mutant. Finally, while Pds5p depletion also shows reduced Smc3p acetylation, the *mcd1-ROCC* allele does not (Chan et al. 2013; Robison, unpublished), again separating the function of this complex conformation in cohesion maintenance from additional functions it may have in promoting acetylation.

A second possibility stems from our observation that the D667 region of the hinge is necessary for the communication between cohesin complexes as revealed by interallelic complementation. We showed that *smc3*-*D667* was unable to complement the inviability of *smc3-42* in *trans*. We previously showed viability of *smc3-42* could be complemented by chromosome bound *smc3-K113R.* Furthermore, the interallelic complementation for viability reflected restoration of all cohesin’s biological functions and restoration of smc3-42p binding to DNA (Eng et al. 2015). Similar phenotypic and molecular interallelic complementation for *mcd1* alleles was also observed (Eng et al. 2015). These observations led us to suggest that interallelic complementation of cohesin mutants reflected cohesin communication likely by the physical interaction between cohesin complexes. The importance of SMC complex oligomerization in their function is gaining traction. The inability of *smc3-D667* to complement *smc3-42* is consistent with the idea that the D667 region of the hinge is necessary for the physical interaction between cohesins and this physical interaction is necessary for maintaining cohesion and condensation.

We propose a working model in which cohesin oligomerizes by forming inverted dimers such that the hinge of one cohesin binds to the head of the other cohesin possibly through binding to Scc3p and that this hinge-head interaction is stabilized by Pds5p. As suggested previously, we can imagine two ways in which hinge-dependent oligomerization might be critical for maintenance of tethering (Eng et al. 2015). We previously showed that mere binding of cohesin to DNA is insufficient to generate tethering, implying that tethering requires an additional activity (Eng et al. 2014). In one model (intramolecular handcuff), two DNA binding activities reside in the same cohesin. In this case oligomerization may inhibit (possibly by physical occlusion) factors that destabilize one of these binding activities. In a second model (intermolecular handcuff) tethering is achieved directly by hinge-dependent oligomerization of two cohesins each of which has a single DNA binding activity. Resolving these models awaits direct biochemical assays for cohesin oligomerization.

## Acknowledgements

We thank Thomas Eng for helpful experimental guidance and the entire Koshland lab for fruitful discussions and reagents. The yeast Smc3-K113 acetylation antibody was a kind gift of Katsuhiko Shirahige. We also thank Benjamin Rowland and Ahmed Elbatsh for advice using the Smc3-K113 acetylation antibody.

## Materials and Methods

### Random insertion screen of *SMC3*

Plasmid pBR25 containing *pGAL-SMC3 URA3 ARS/CEN* was subject to *in vitro* transposition according to the protocol recommended by the MuA transposase MGS Kit (ThermoFisher Cat. F701). After transforming into TOP10 cells (Thermo), 5,756 AmpR KanR colonies were pooled and plasmids harvested by Midi Prep (Qiagen). The pooled library was digested with NotI to excise the KanR marker, gel extracted, and religated. Ligation products were transformed once again into TOP10 cells and confirmed to have lost KanR by replica plating. > 30,000 colonies were pooled, and plasmids harvested by Midi Prep to obtain a library of *pGAL-SMC3* plasmids with fifteen extra nucleotides randomly inserted. Library depth was calculated by multiplying the fraction of pBR25 coding for *SMC3* (3,693 bp of 10,083 bp total) by the number of AmpR KanR colonies (5,756) to obtain 2,118 plasmids expected to have an insertion in *SMC3*. From this calculation, we expect plasmids represented in the library harboring insertions every approximately 1.7 base pairs along *SMC3*. The library was transformed into wild-type (3349-1B) and *smc3-42* (3358-3B) strains which were incubated at 23°C for three days to select for transformants on synthetic complete media lacking uracil (SC –URA) with 2% dextrose supplied as the carbon source. 3,382 wild-type colonies and 1,811 *smc-42* colonies were screened. Transformation colonies were replica plated onto SC –URA 2% galactose plates and SC –URA 2% dextrose plates as a control and incubated overnight at 23°C. Colonies that were slow growing or inviable on galactose plates were then grown overnight in liquid YPD and plated in 10-fold serial dilutions on 1) galactose plates to confirm slow growth and 2) 5-FOA plates with 2% galactose to confirm linkage of slow growth to presence of the RID library plasmid. Insertion mutations were identified by PCR and sequencing across the entire *SMC3* ORF.

### Yeast strains, media, and growth

All strains used are in the A364A background and their genotypes can be found in the Strain List. Yeast extract/peptone/dextrose media and synthetic dropout media was prepared as previously described (Guacci et al. 1997). Conditional AID degron strains were grown in YPD and auxin (3-indoleacetic acid, Sigma Aldrich Cat I3750) added to a final concentration of 0.75 mM to deplete AID-tagged proteins. YPD agar plates supplemented with auxin were made by cooling molten YPD 2% agar to 55°C prior to addition of auxin.

### Cohesion assays

Sister chromatid cohesion was assessed at either the centromere-distal *LYS4* locus or centromere-proximal *TRP1* locus on chromosome IV in which *LacO* arrays had been integrated. The *GFP-LacI* fusion allele integrated at *HIS3* allows fluorescence microscopic visualization of *LacO* arrays. Cohesion was scored by growing cells to mid-log phase (OD600 ∼0.3) and arresting them in G1 using alpha factor at 10_-8_ M (Sigma Aldrich). After arresting for 3 hours, auxin was added to a final concentration of 0.75 mM to deplete Smc3-AIDp for one hour. Cells were released from G1 arrest by washing in YPD containing auxin and 0.1 mg/mL Pronase E (Sigma Aldrich) five times and resuspending in YPD containing auxin and 15 μg/mL nocodozole (Sigma Aldrich). Cultures were incubated at 23°C and samples fixed either 1) periodically for assessing S-phase cohesion establishment or 2) after three hours in which > 95% of cells had arrested in G2/M. In addition to fixation for microscopy, samples were taken in parallel to assess DNA content by flow cytometry. Cohesion was scored by counting the number of GFP-LacI foci in the nucleus by fluorescence microscopy of fixed cells.

### Monitoring condensation at the *rDNA* locus

Cells were grown as if for assessing cohesion by arresting in YPD containing auxin and nocodazole following release from G1. Cells were fixed, spheroplasted, and lysed to allow binding of chromosomes to slides as described previously (Guacci et al. 1994). Briefly, 1 mL of mid-M phase arrested cells were fixed two hours in 100 uL of 37% formaldehyde, washed twice in water, and spheroplasted for one hour. Triton X-100 was added to 0.5% for 5 minutes, then cells were pelleted and resuspended in water. Cells were then added to poly-lysine-coated slides for ten minutes. 0.5% SDS was added for 10 minutes to solubilize membranes and release DNA masses then removed. Slides were fixed in 3:1 methanol:acetic acid for five minutes and allowed to dry. Cells on slides were treated with RNase A and Proteinase K and subject to a series of short 70%, 80%, 90%, and 100% ethanol washes. After drying, DNA masses were visualized with DAPI and *rDNA* morphology scored.

### Chromatin immunoprecipitation (ChIP)

Cells were grown as if for assessing cohesion by arresting at mid-M phase in YPD containing auxin and nocodazole following release from G1 arrest. ChIP was performed as described previously (Eng et al. 2014; Wahba et al. 2013) except that chromatin shearing was performed on a Bioruptor Pico machine (Diagenode, Denville, NJ) for 5 minutes (30 sec on/off cycling). Immunoprecipitation was performed using monoclonal Mouse anti-HA (Roche), monoclonal Mouse anti-V5 (ThermoFisher), polyclonal Rabbit anti-Pds5p (Covance Biosciences, Princeton, NJ), or polyclonal Rabbit anti-Mcd1p (Covance) antibodies. A no antibody control was always included to assess specificity of chromatin recovery.

### Detection of Smc3-K113 acetylation by Western blotting

Cells were grown to OD_600_=0.5 in YPD at 23°C before addition of auxin to 0.75 mM and incubation for 1 hour. Nocodazole was added to a final concentration of 15 μg/mL to arrest cells in mid-M phase. Cells were pelleted and resuspended in lysis buffer consisting of 25 mM HEPES pH 8.0, 2 mM MgCl_2_, 100 μM EDTA, 500 μM EGTA, 1% NP-40, 150 mM KCl, 15% glycerol, Complete-Mini EDTA-free protease inhibitor cocktail (Roche), 10 mM sodium butyrate, and 20 mM beta-glycerophosphate. Cells were incubated in buffer for 30 minutes on ice, then glass beads were added to a 1:1 volume ratio before bead-beating for three minutes. Lysates were pelleted at 14K for 10 minutes at 4°C, and protein concentration measured using Coomassie Brilliant Blue. Lysates were boiled in 120 mM HEPES pH 7.0 containing 1% SDS at 95°C for five minutes, then diluted 1:1 in 2X Laemmli sample buffer. Smc3-K113 acetylation was detected by blotting with monoclonal Mouse antibody (a gift from K. Shirahige) at a concentration of 1:1,000 in 5% milk-PBST.

### Chromosome spreads and microscopy

Cells were grown as if for assessing cohesion by arresting in mid-M phase in YPD containing auxin and nocodazole following release from G1 arrest. Chromosome spreads were prepared as described previously (Wahba et al. 2013). Slides were incubated with 1:5,000 rabbit polyclonal anti-Mcd1p and 1:5,000 mouse anti-V5 antibody (Life Technologies). Antibodies were diluted in blocking buffer (5% BSA, 0.2% milk, 1X PBS, 0.2% Triton X-100). Secondary Alexa Fluor 488-congugated chicken anti-mouse and Alexa Fluor 568-congugated donkey anti-rabbit (ThermoFisher Cats. A21200 and A10042) antibodies were diluted 1:5,000 in blocking buffer. Indirect immunofluorescence was detected on an Axioplan2 microscope (Zeiss, Thornwood, NY) using the 100X objective (numerical aperture 1.40) which is equipped with a Quantix charge-coupled camera (Photometrics).

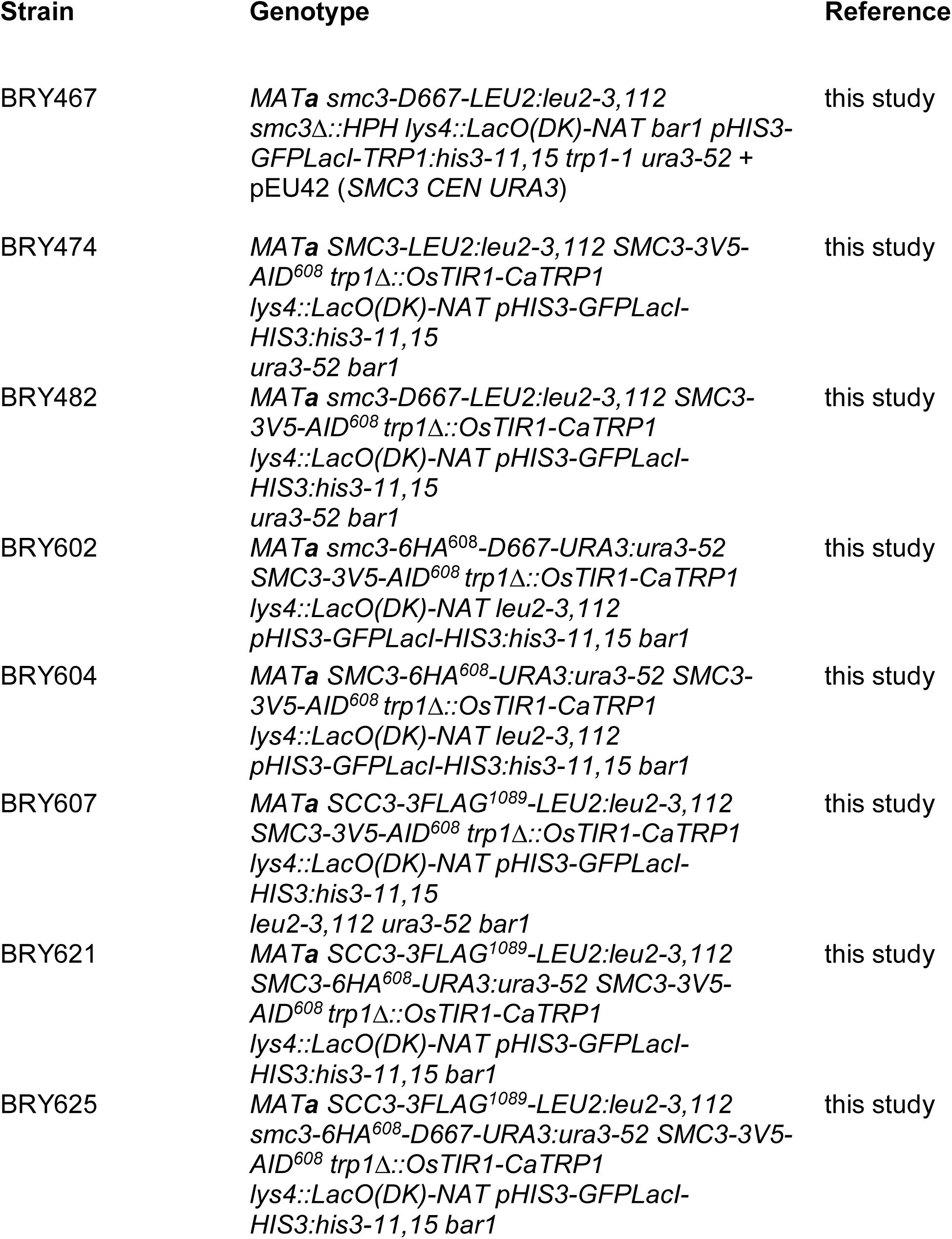
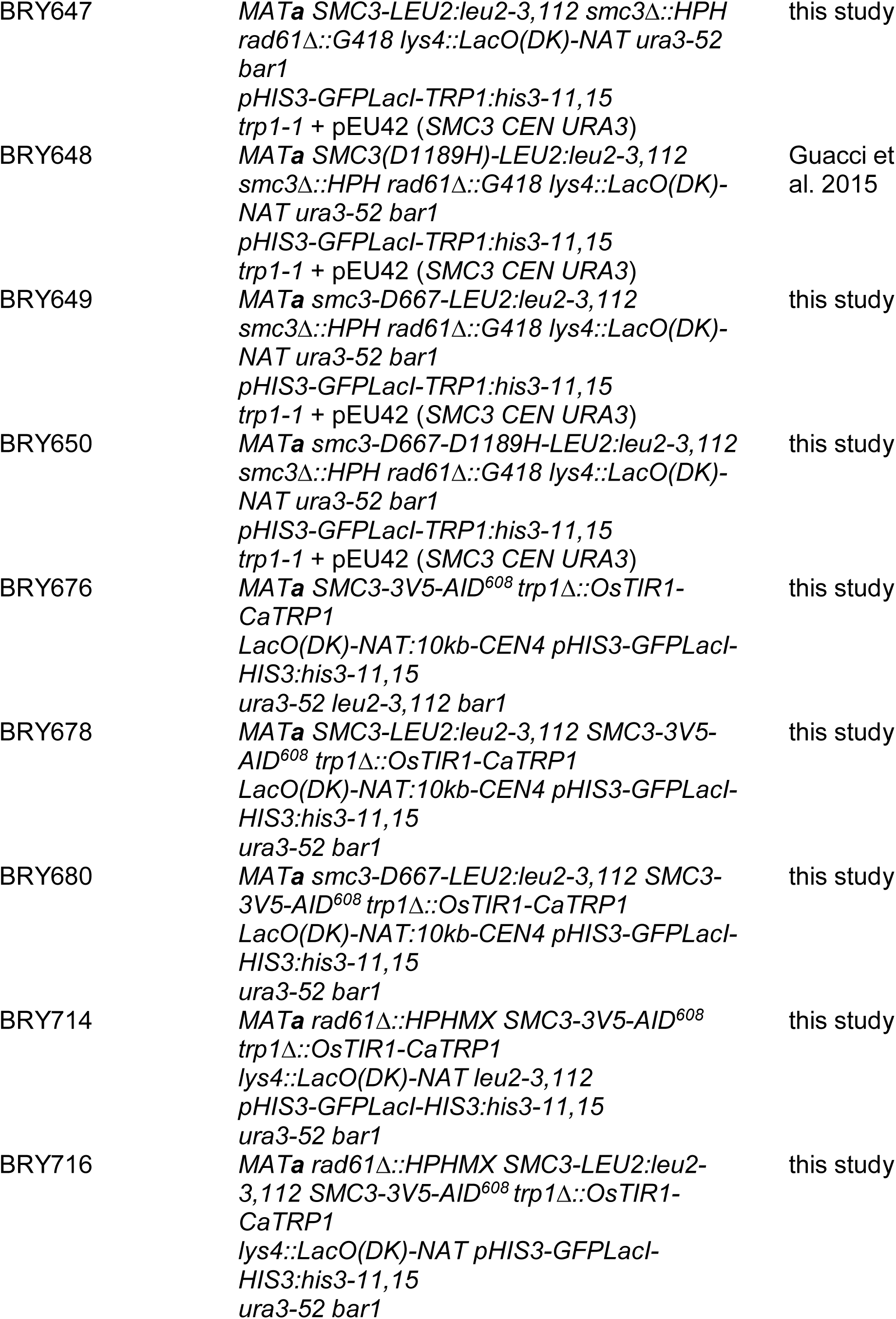
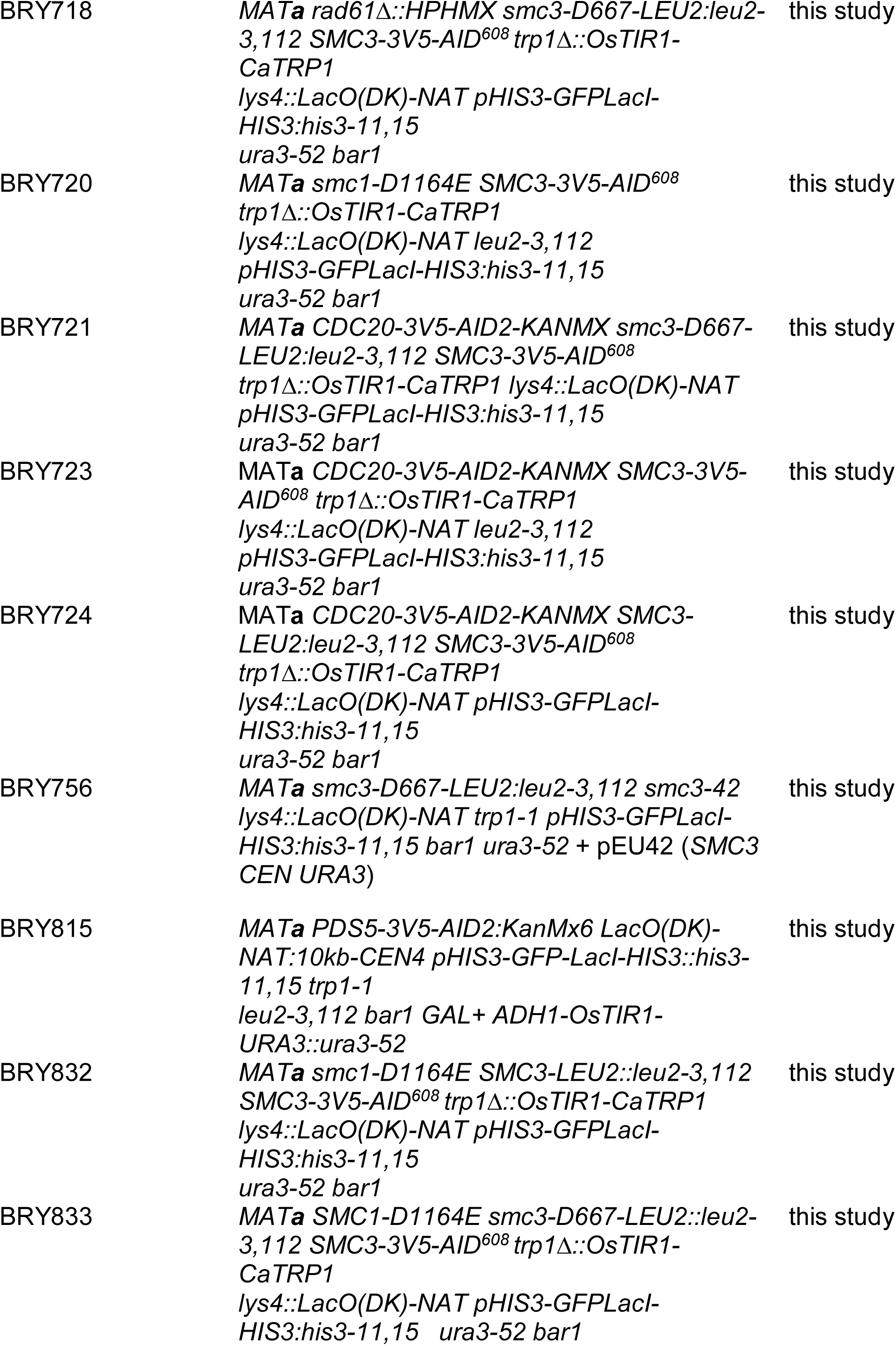
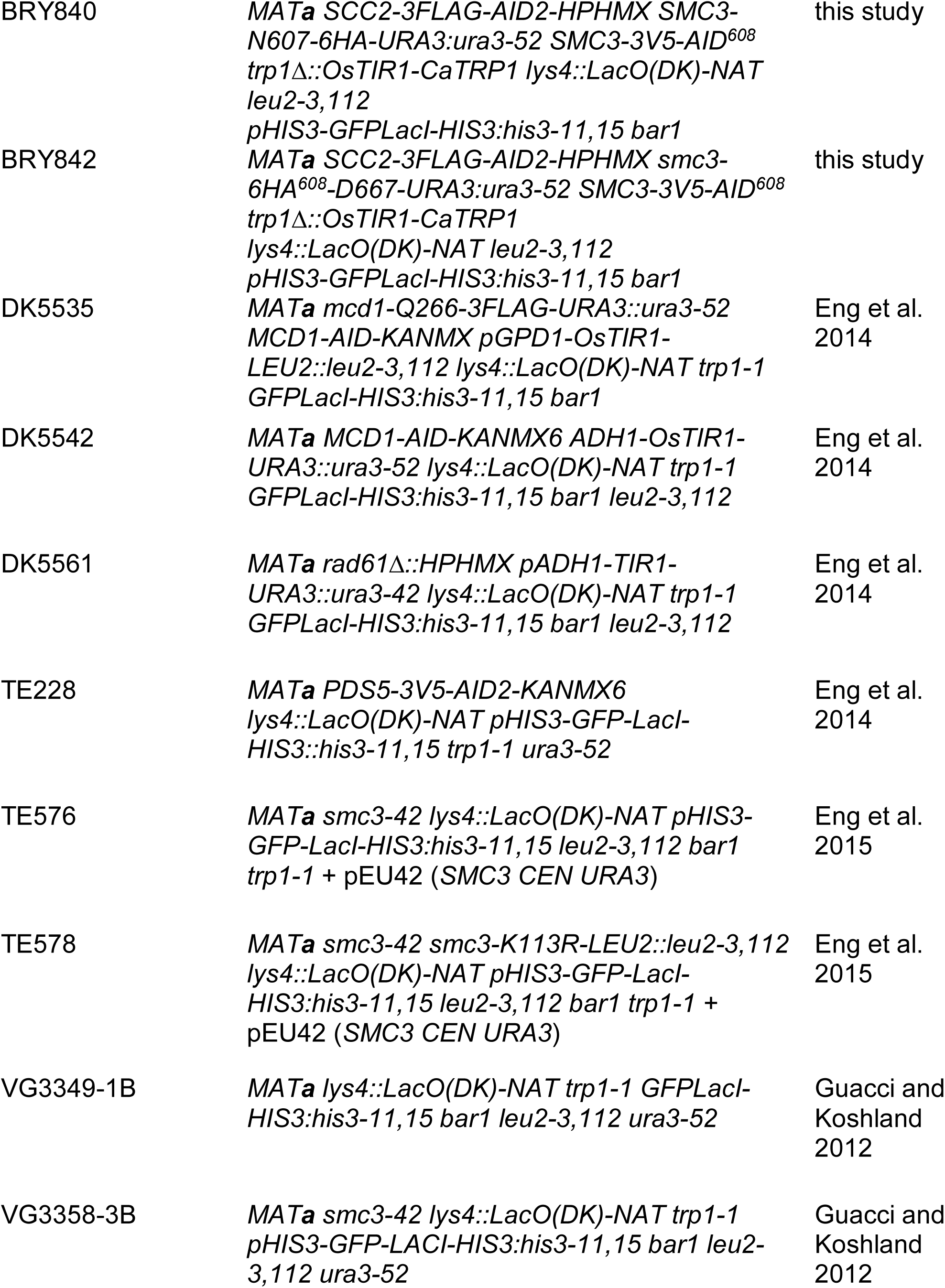
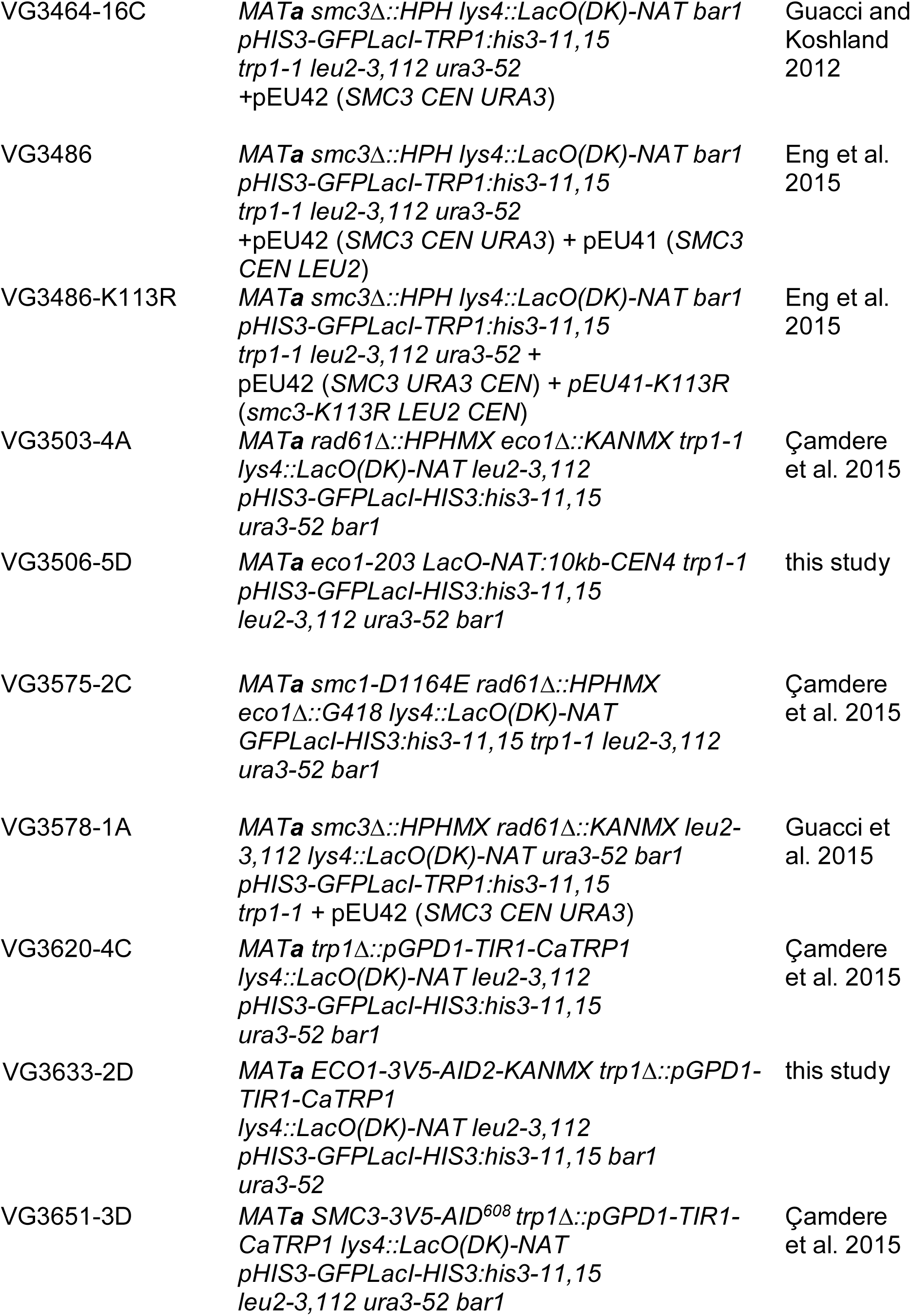

## Figure Legends

**Supplementary Figure 1.**
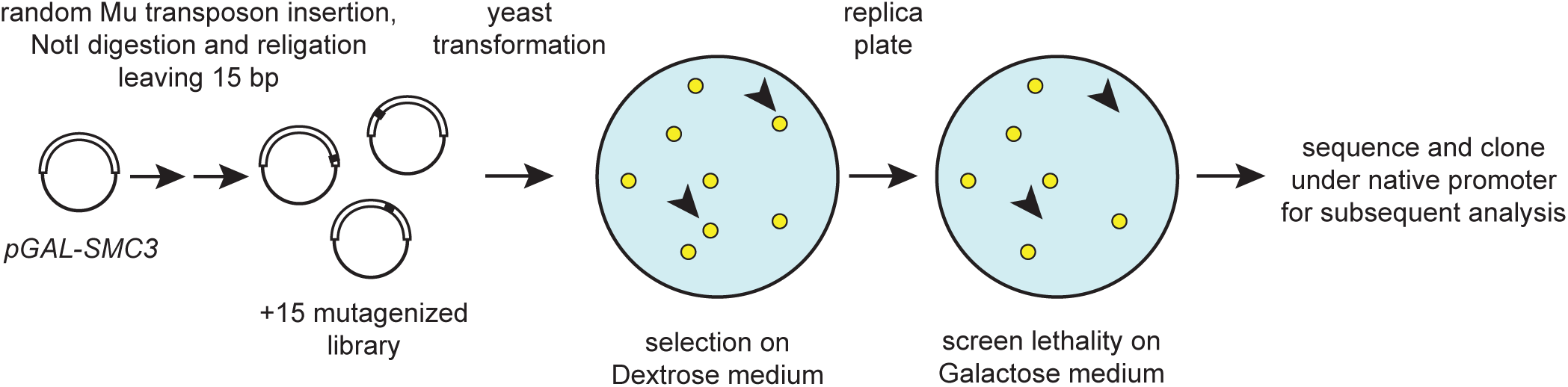
*SMC3* random insertion dominant (RID) screen workflow. A *pGAL-SMC3* URA3 CEN/ARS plasmid, pBR25, was subject to *in vitro* transposase mutagenesis to generate the RID library which consists of plasmids with fifteen additional nucleotides randomly inserted (see Materials and Methods). Haploid yeast were transformed with the SMC3 RID library and selected on dextrose plates. Transformants were replica plated to galactose plates to induce expression by *pGAL*. Mutants that were inviable or had slow growth on galactose were tested to confirm that the RID plasmid was the cause of this phenotype. Confirmed RID plasmids were sequenced to determine insertion location.

**Supplementary Figure 2.**
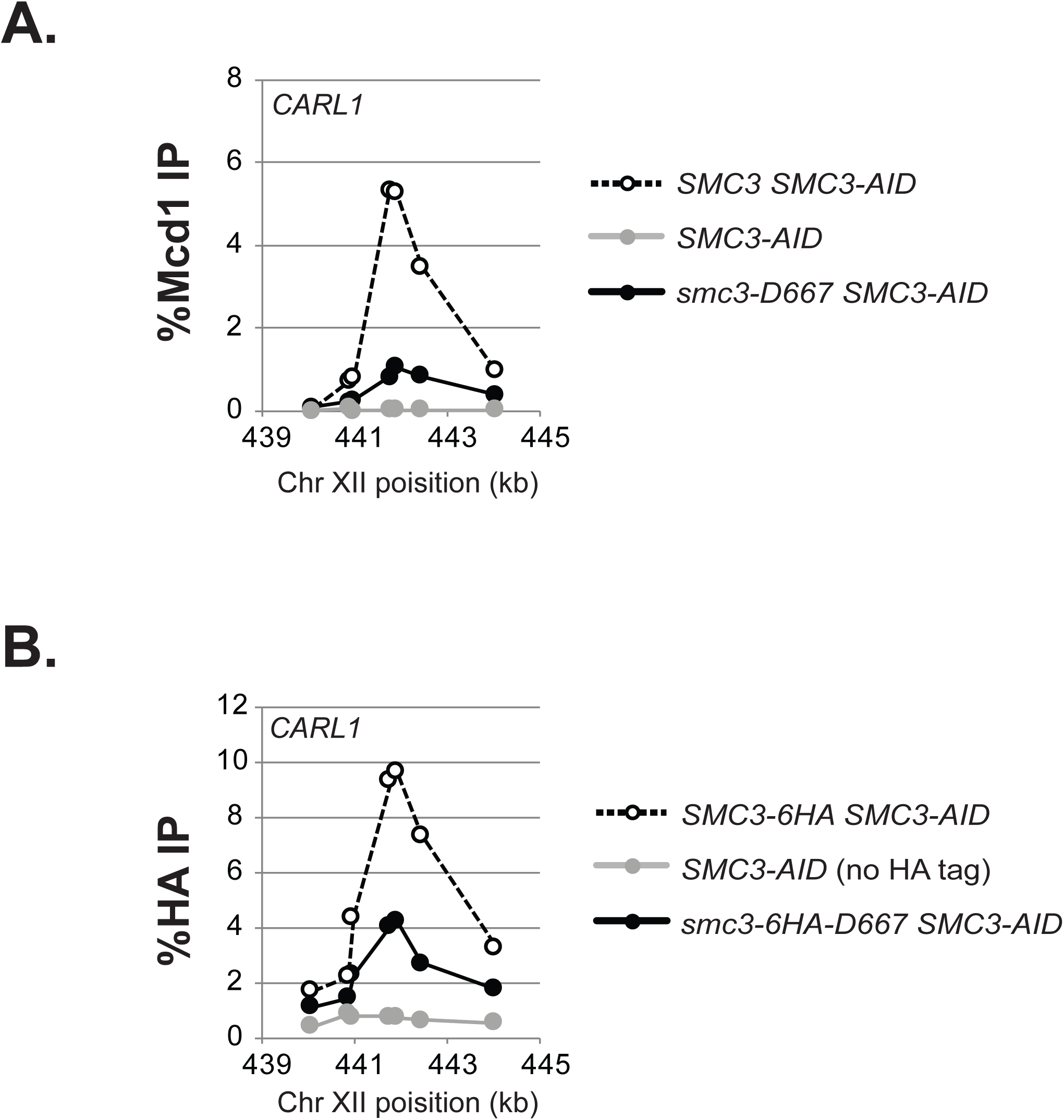
Assessment of smc3-D667p cohesin binding to *CARL1*. A. ChIP of Mcd1p binding at the *CARL* locus. Samples from Figure 2C assayed for Mcd1p binding to *CARL.* Wild-type strain *SMC3 SMC3-AID* (dotted line), *smc3-D667 SMC3-AID* strain (black line) and *SMC3-AID* alone (grey line). B. ChIP of HA epitope tagged Smc3p and smc3-D667p at the *CARL* locus. Samples from Figure 2D assayed for Smc3p and smc3-D667p binding to *CARL. SMC3-6HA SMC3-AID* (dotted line), *smc3-6HA-D667 SMC3-AID* (black line) and *SMC3-AID* only (grey line).

**Supplementary Figure 3.**
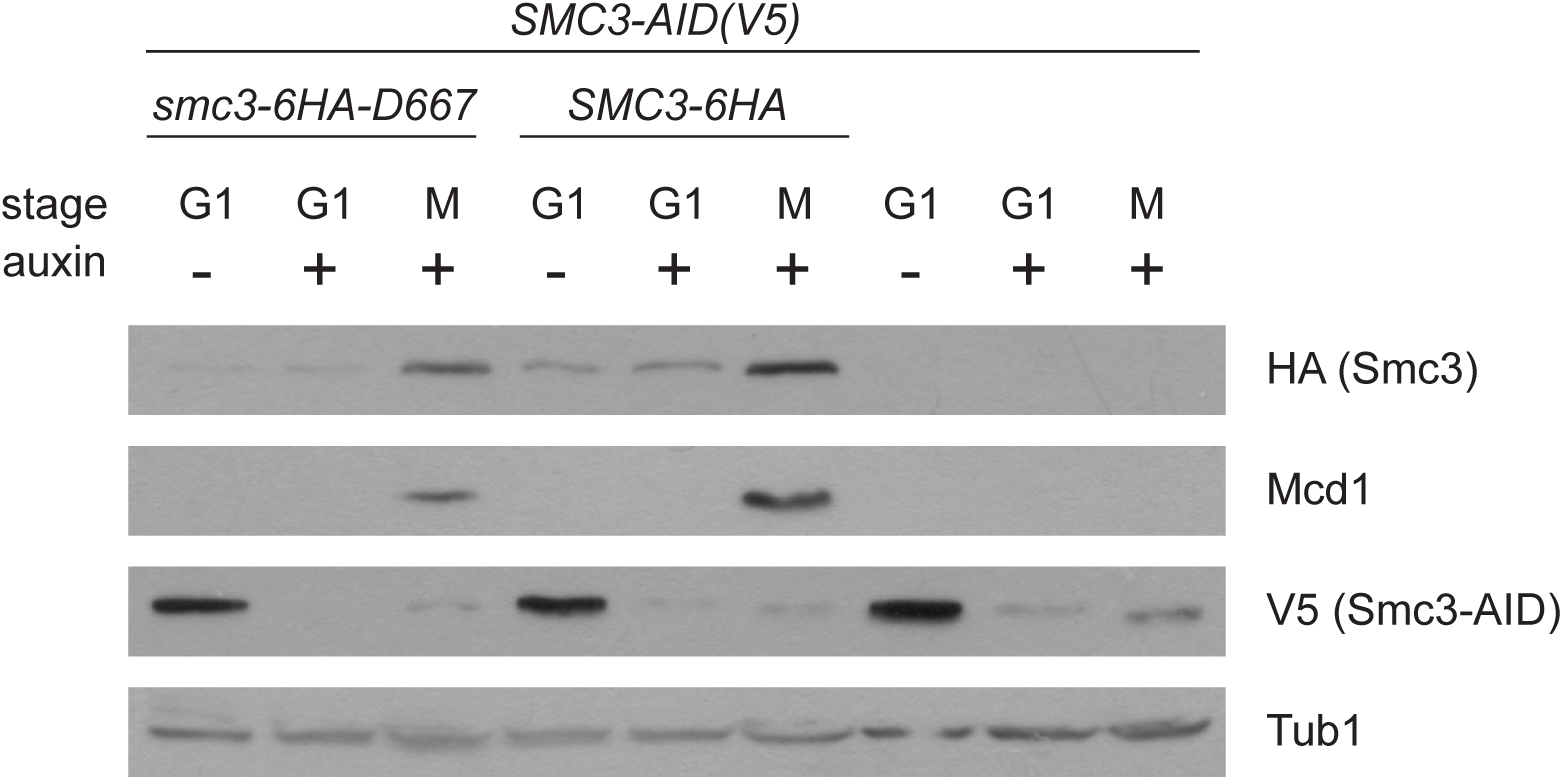
Western analysis showing depletion of Smc3-AIDp and levels of cohesin subunits Mcd1p and HA-tagged Smc3p. Protein extracts from *SMC3-3V5-AID* strains expressing *SMC3-6HA*_*607*_*-D667* (BRY602), *SMC3-6HA*_*607*_ (BRY604), or no additional *SMC3* allele (VG3561-3D) in Figure 2D.

**Supplementary Figure 4.**
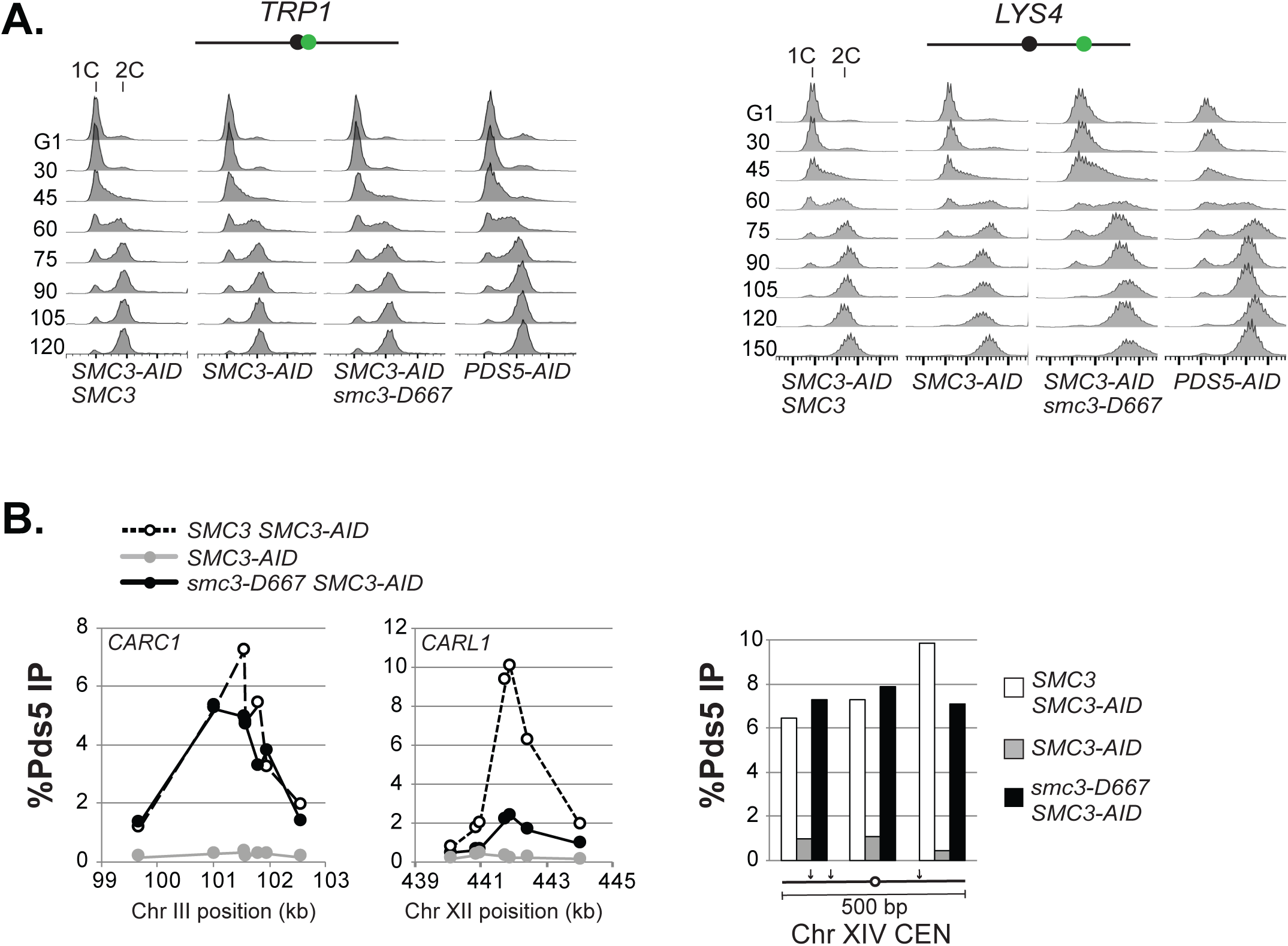
A. Flow cytometry to assess cell-cycle progression of cells from experiments in Figure 3C. Strains bearing the LacO array near the *CEN*-proximal *TRP1* locus (left) and *CEN*-distal *LYS4* locus (right). B. ChIP of Pds5p binding at the *CEN*-proximal *CARC1* and *CEN*-distal *CARL* loci (left) and *CEN14* (right). ChIP of samples from Figure 3D showing Pds5p binding to *CARC1, CARL*, and *CEN14.* Wild-type strain *SMC3* (dotted lines and white bars), *smc3-D667* strain (black lines and black bars) and *SMC3-AID* alone (grey lines and grey bars).

**Supplementary Figure 5.**
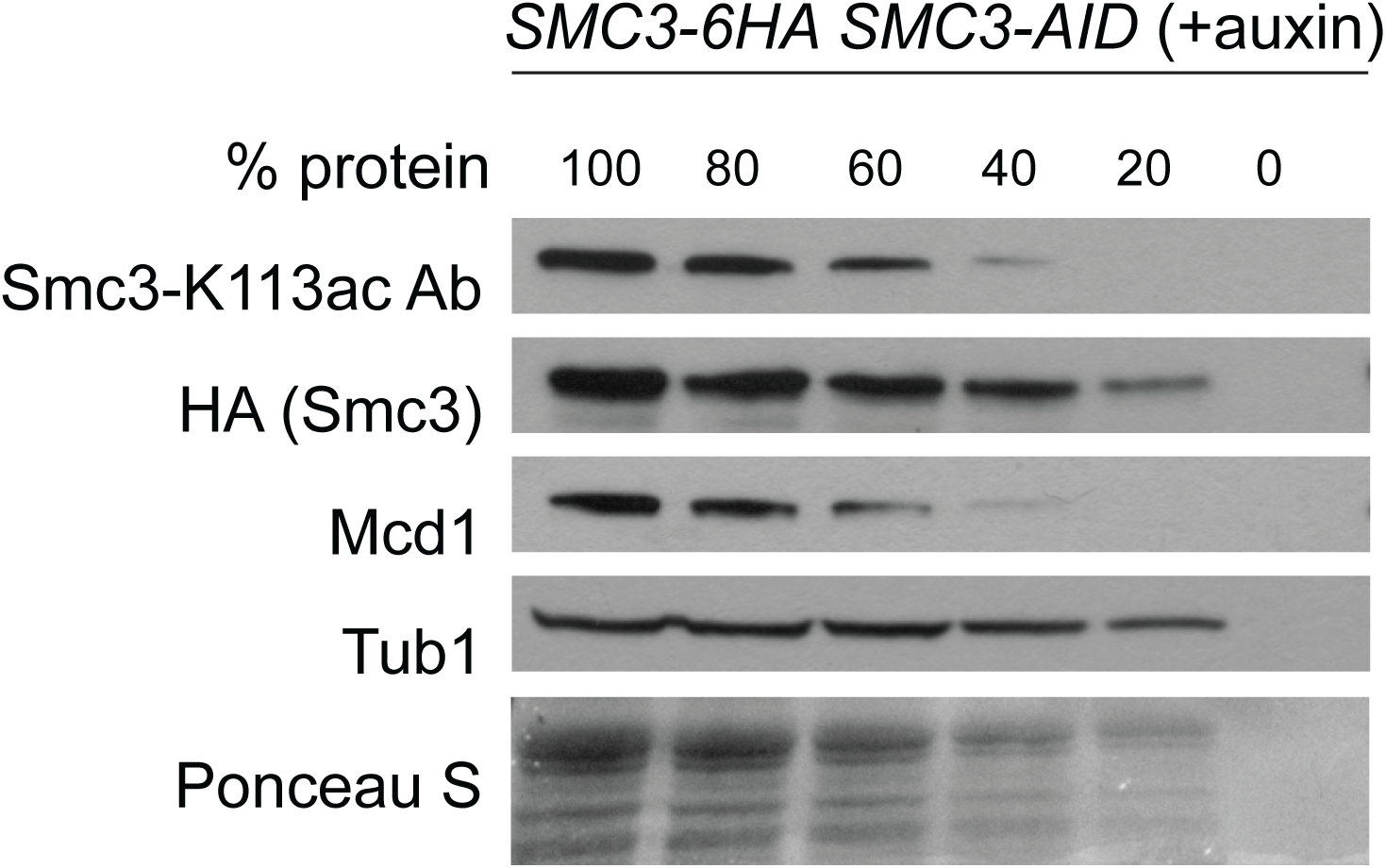
Non-linearity of acetylated Smc3-K113 specific antibody, related to Figure 5. A culture of the wild-type *SMC3-6HA SMC3-AID* (BRY604) haploid strain was grown as described in Figure 5A. Total protein extract from mid-M phase-arrested cells was obtained as described in Materials and methods. Extract was diluted 1:2 in buffer containing 120mM HEPES pH 7.0 and 1% SDS and boiled 5 minutes at 95%. Boiled extract was then diluted 1:2 in 2X Laemmli sample buffer to create the 100% protein sample. This sample was then diluted in 2X Laemmli buffer to 80%, 60%, 40% and 20% concentration and subjected to SDS-PAGE and Western analysis using the indicated antibodies.

**Supplementary Table 1.**

| Location | Insertion | Viability 5-FOA 23°C |
| --- | --- | --- |
| D84 | CGRND | Not Tested (NT) |
| D127 | AAAGD | - |
| P147 | LRPQP | - |
| L165 | RPQQL | + |
| G171 | AAAAG | - |
| N204 | AAALN | - |
| Y253 | NAAAY | + |
| S343 | IAAAS | - |
| N517 | RPQAN | + |
| D643 | CGRKD | + |
| K1023 | VRPHK | - |
| Y1164 | CGRKY | NT |

**Supplementary Table 2.**
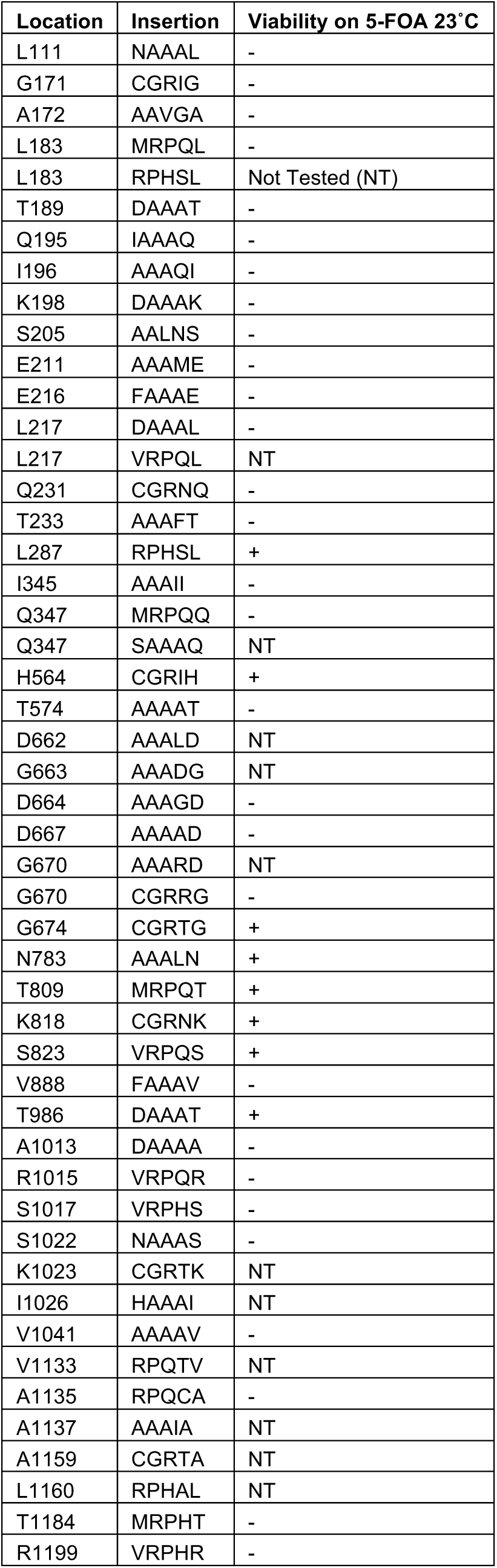

